# Inferring cellular and molecular processes in single-cell data with non-negative matrix factorization using Python, R, and GenePattern Notebook implementations of CoGAPS

**DOI:** 10.1101/2022.07.09.499398

**Authors:** Jeanette Johnson, Ashley Tsang, Jacob T. Mitchell, Emily Davis-Marcisak, Thomas Sherman, Ted Liefeld, Melanie Loth, Loyal A Goff, Jacquelyn Zimmerman, Ben Kinny-Köster, Elizabeth Jaffee, Pablo Tamayo, Jill P. Mesirov, Michael Reich, Elana J. Fertig, Genevieve L. Stein-O’Brien

## Abstract

Non-negative matrix factorization (NMF) is an unsupervised learning method well suited to high-throughput biology. Still, inferring biological processes requires additional post hoc statistics and annotation for interpretation of features learned from software packages developed for NMF implementation. Here, we aim to introduce a suite of computational tools that implement NMF and provide methods for accurate, clear biological interpretation and analysis. A generalized discussion of NMF covering its benefits, limitations, and open questions in the field is followed by three vignettes for the Bayesian NMF algorithm CoGAPS (Coordinated Gene Activity across Pattern Subsets). Each vignette will demonstrate NMF analysis to quantify cell state transitions in public domain single-cell RNA-sequencing (scRNA-seq) data of malignant epithelial cells in 25 pancreatic ductal adenocarcinoma (PDAC) tumors and 11 control samples. The first uses PyCoGAPS, our new Python interface for CoGAPS that we developed to enhance runtime of Bayesian NMF for large datasets. The second vignette steps through the same analysis using our R CoGAPS interface, and the third introduces two new cloud-based, plug-and-play options for running CoGAPS using GenePattern Notebook and Docker. By providing Python support, cloud-based computing options, and relevant example workflows, we facilitate user-friendly interpretation and implementation of NMF for single-cell analyses.

## Introduction

The central challenge of high throughput biology, as exemplified by single-cell analysis, pertains to the reduction of extremely high-dimensional data into a format from which humans can observe patterns, formulate mechanistic hypotheses, and design new experiments. High-throughput experiments are ubiquitous across many areas of biomedical and biological research. As technology evolves to perform these experiments, algorithmic strategies and computing capabilities must evolve just as swiftly to keep up with the sheer amount of data they yield. Non-negative matrix factorization (NMF) is a mathematical technique with a long history in the field of genomics for the analysis of bulk RNA sequencing data^1,2^, and it has been widely adopted as a powerful dimensionality reduction tool for single-cell data as well^3–10^. As many phenotypes are unknown *a priori* in single-cell data, this learning method is particularly well suited for unsupervised analyses. The additive nature of solutions from NMF yields interpretable patterns that can be associated directly with biological processes. Thus, NMF solutions, by definition, encode many characteristics of each cell simultaneously, including identity, state transitions, molecular processes, and even technical artifacts^4,6,9–17^.

Multiple software packages implement non-negative matrix factorization, many of which apply to single-cell data^18–22^. Still, biological interpretation of NMF solutions requires further functionalization and practical, end-to-end workflows developed specifically for omics data. Technical components, such as algorithm assumptions, convergence, and dimensionality all impact the analysis findings^23^. Biologically interpretable solutions of NMF analysis also rely on custom, post hoc visualization and statistics of the patterns learned from the data^24^. These steps are often customized for each analysis and are not previously codified into a cohesive description of the workflow required for interpretable NMF analysis.

Here we present three protocols for interpretable analysis of single-cell RNA-seq (scRNA-seq) data with our sparse, Bayesian NMF algorithm Coordinated Gene Activity in Pattern Sets (CoGAPS^19^) based on previous findings of its robustness to initial conditions^4,12,13^. CoGAPS was originally released in an R/Bioconductor package by the same name^19,25^. This protocol presents a new Python interface for CoGAPS, called PyCoGAPS, to enhance accessibility of this method. Our three protocols then demonstrate step-by-step NMF analysis across distinct software platforms. They are applied to characterize malignant epithelial cell state transitions in pancreatic cancer using published single-cell RNAseq data from 15,219 epithelial cells from 42 tumor and 11 control samples, as we annotated previously^26^. The first protocol uses our new Python interface, which we have found to perform faster for larger matrices. The second protocol uses the original R interface for CoGAPS, but emphasizes optimal practices for single-cell data. The third protocol demonstrates two options for running CoGAPS with large single-cell RNA-Seq datasets using pre-defined, cloud-based computing environments built with GenePattern Notebook^27^ and Docker. This range of options makes NMF accessible to users regardless of their programming background or access to computing architecture.

## Overview of NMF algorithms for single-cell analysis

NMF approximates an input data matrix as the product of two lower-dimensional matrices with non-negative entries. If the input matrix of single-cell data contains genes along its rows and cells along its columns, the first result matrix is of dimension genes-by-patterns and the second patterns-by-cells. The number of patterns (or equivalently, features) that define the inner dimension of the two matrices in the factorization is an input variable to the algorithm, which will here be referred to as *k*, which is represented in our code as parameter *nPatterns*. When applying NMF to analyses of other high-dimensional data modalities, “genes” and “cells” in this protocol could be replaced by any number of other variables, depending on the experiment and measurement technology. Following the standardized notation for factorization analyses from Stein-O’Brien et al^24^, we here refer to the genes-by-patterns matrix as the *amplitude matrix* (**A**) and the patterns-by-cells matrix as the *pattern matrix* (**P**). A variety of alternate nomenclature has been assigned to these matrices in other studies; often the amplitude matrix is referred to as the weights matrix^28^ or meta-genes^1^ and the pattern matrix as the heights matrix^28^ or meta-cells. The non-negativity assumption in NMF yields nonnegative features in these matrices that add together to reconstruct the signal in the input data (Fig 1). This non-negative constraint contributes to the solution’s biological interpretability, as negative quantities do not exist in nature^24^. Together, the amplitude matrix describes the association of each gene with each pattern, and the pattern matrix provides information about the relative contribution of each pattern to the phenotype of each cell or sample.

**Figure 1.**
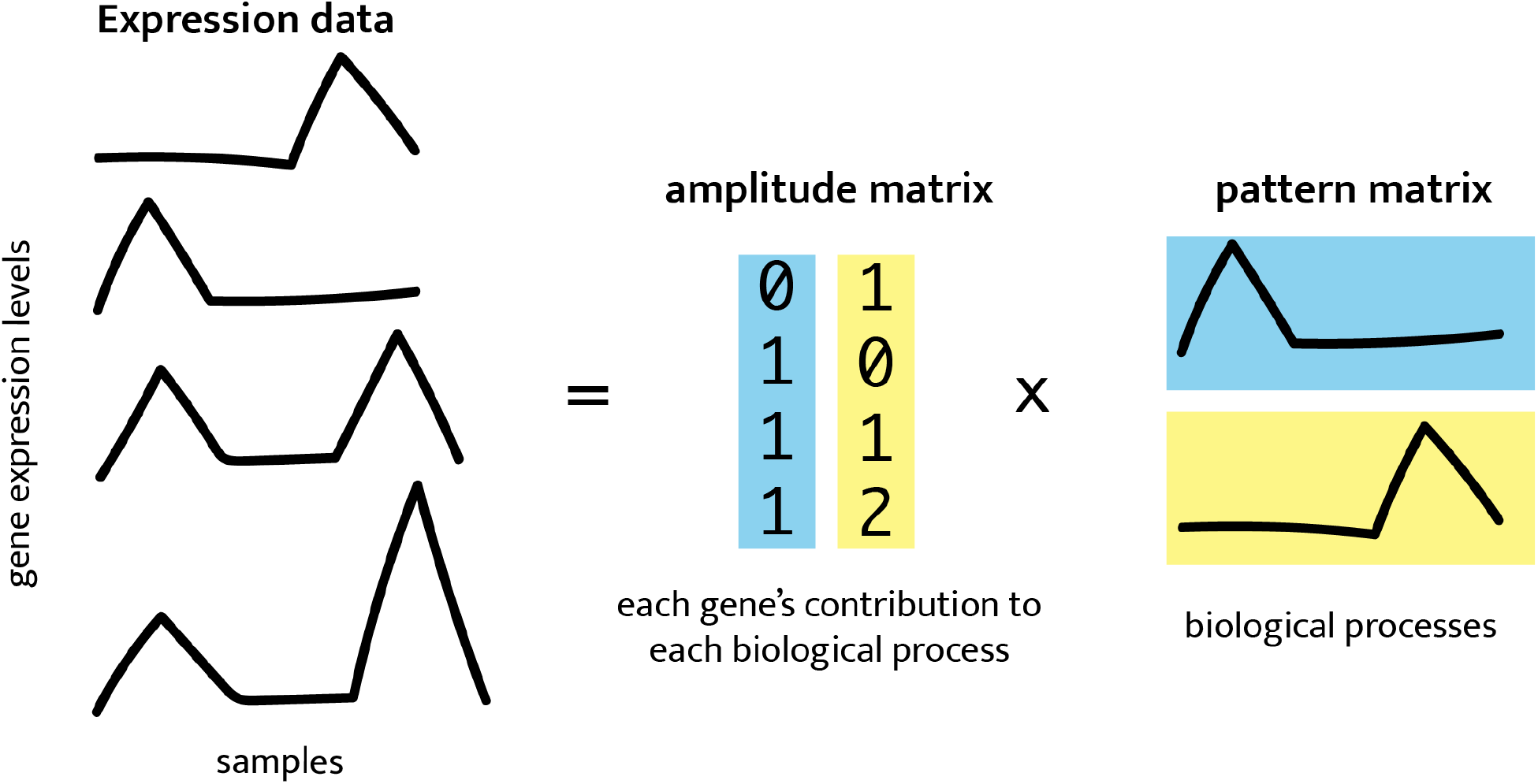
NMF factorizes expression data into lower-dimensional amplitude (A) gene weights matrix and pattern (P) weights matrix whose product approximates the input.

There are a wide variety of NMF techniques used for high throughput molecular analysis, most recently scRNA-seq analysis^4,6,9–16^. Algorithms used to solve the NMF problem can be divided into two major classes: **gradient-based** methods that seek a single solution that optimizes a cost function^28–30^ and **Bayesian methods** that estimate the posterior distribution of the amplitude and pattern matrices^8,21,31^. Both classes can be modified to encode additional constraints on top of non-negativity, further differentiating the various NMF techniques. For example, the Bayesian NMF CoGAPS^3,19,25^ and gradient-based LS-NMF^32^ both model the uncertainty in the expression data in the factorization. In addition, CoGAPS also leverages the Bayesian architecture through an atomic prior^33^ to model sparsity in both the amplitude and pattern matrices. The three protocols presented here are intended for use with CoGAPS. However, all of these protocols are readily adaptable for analysis with other NMF algorithms or even other forms of matrix factorization.

## Key considerations for NMF analysis

A generalized workflow for NMF analysis of single-cell data is summarized in Fig 2. Each step in this workflow is described generally to facilitate customization of the template protocols to other factorization methods, non-negative and otherwise. This section discusses several best practices and open questions for NMF, and offers strategies for choosing parameters and assessing the learned solutions.

**Figure 2:**
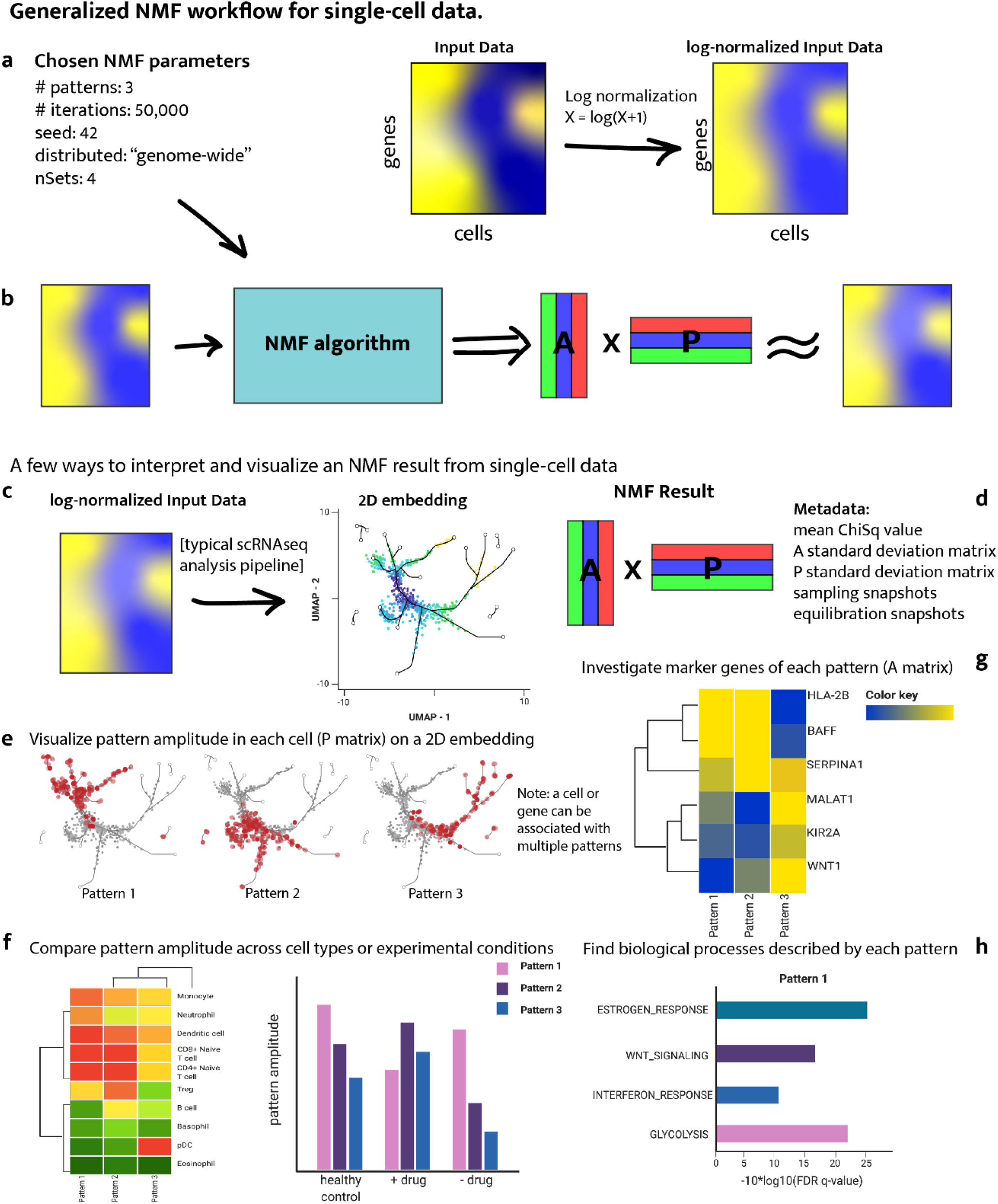
A generalized workflow for performing NMF on single-cell data. (a) NMF algorithms take as input a list of parameters and a data matrix. For scRNA-seq data, the counts matrix should be log-normalized. (b) NMF yields an amplitude matrix (A) and a pattern matrix (P) which approximately factorize the input data. (c) NMF results can supplement a dimension reduction analysis pipeline. (d) An NMF result typically consists of A and P matrices along with metadata about the run. (e) To visualize the pattern weight in each cell, the P matrix can be used to color a UMAP or other dimension-reduction plot. (f) The P matrix can also be used to compare pattern weights across cell types or experimental conditions. (g) The A matrix can be used to find marker genes for each pattern, which can then be useful in (h) gene set enrichment analysis, identifying biological processes and terms associated with each pattern.

### Data preprocessing and input

The majority of NMF analyses are performed on normalized and log transformed data^34,35^, which is recommended as a preprocessing step in our CoGAPS protocols. We note that regardless of how the input data is transformed, it must contain only non-negative values, as this is a central requirement of NMF. We note that some emerging NMF algorithms have error models designed for raw count data^36^, and therefore do not require this normalization.

Many scRNA-seq technologies are subject to drop-out, resulting in zero values for a large proportion of measurements from technical rather than biological conditions. Several imputation approaches have been developed to estimate the signal in these missing data prior to analysis^37^. Still, it is not necessary to impute the input data for NMF analysis and indeed the reconstruction of the data estimated from the product of the inferred amplitude and pattern matrices can be used as an alternative imputation scheme^38^. Moreover, the sparsity model in the atomic prior from CoGAPS is tailored to the sparsity of scRNA-seq data motivating our selection of this algorithm as the foundation for this protocol^3,25^. If the user desires, imputed data is acceptable for input, but we note that the imputation algorithm employed will impact the inferred solutions.

Technical aspects of scRNA-seq experiments, such as library, processing day, dissociation quality, etc, can introduce further artifacts in the signal from scRNA-seq data leading to numerous batch correction approaches for scRNA-seq data^39^. Some batch correction algorithms do not change the raw data and focus instead on aligning the embedding used to visualize scRNA-seq data^40,41^, and therefore would not impact the factorization results. Other batch correction algorithms attempt to remove these technical signals from the data^42^. These batch correction approaches may also affect the solution and should be used with caution. Especially as some algorithms, such as CoGAPS, have been demonstrated to concurrently learn technical and biological signals making preprocessing to eliminate batch effects unnecessary^3,4^. Likewise, NMF approaches can also provide a unified embedding between datasets^10^. We acknowledge that these first steps must often vary greatly depending on the biological context, and invite the user to validate optimized custom preprocessing workflows for that context. Comprehensive reviews of preprocessing pipelines for scRNA-seq data have been previously published^43–45^.

### Iterative assessment of optimality of solutions

Biological inference based upon solutions of the amplitude and pattern matrices for a dataset relies on the assumption that the NMF algorithm has returned a stable and biologically relevant factorization. Determining optimality of factorization remains an open question, with various metrics developed to assay performance. These metrics will vary based on the type of NMF analysis used. Bayesian methods for NMF, including CoGAPS, estimate the posterior probability distribution for amplitude and pattern matrices. Bayesian NMF methods for genomics analysis employ a wide variety of Markov chain Monte Carlo (MCMC) and variational algorithms to learn these distributions. Whereas gradient-based and variation methods are subject to local minima, many MCMC methods are designed to overcome local optima, which is crucial in biological applications where there may be many semistable states and thus many local optima. However, this gain in the global optimality of solutions occurs at a cost: these algorithms must be run over many iterations, often resulting in long runtimes, which can be addressed with parallelization^12,25,46^ or GPU computing^21^. Likewise, the local optima of gradient-based techniques can be overcome by leveraging parallel computing to determine the global optima by sampling solutions from multiple initial conditions.

After an MCMC run on a given dataset is complete, it remains to be assessed whether it was run for a sufficient number of iterations to attain accurate sampling from the posterior distributions for both the amplitude and pattern matrices — a property known as convergence — and whether the user-specified number of patterns learned corresponds appropriately to the biological question under investigation. When convergence is reached, increasing the number of iterations will enhance the density of sampling from the posterior distribution to improve analytic estimates of the distribution but will no longer improve the learned solution. Estimating the optimal number of iterations requires some shrewdness on the part of the user to decide how many iterations are required to produce stable and reasonable results for their data or application. The application of Bayesian convergence metrics to determine the stopping criterion for Bayesian NMF algorithms remains an open area of research. Therefore, it is critical to evaluate the stability of the likelihood calculation over the chain to assess the optimal number of iterations for each Bayesian NMF algorithm.

The convergence metrics for each NMF algorithm depends on the details of the mathematical formulation of the model used for the factorization. In the case of CoGAPS, this algorithm performs factorization of a transcriptional dataset *D*_*i,j*_ with genes (*i*) and cells*(j)*. according to the Bayesian model 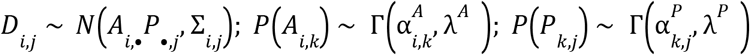 where *N*(•, •) indicates a univariate normal distribution, the shape parameters are modeled according to a Poisson prior with hyperparameter α, and the additional hyperparameters are fixed to model transcriptional data^3,46^. Implementing this model through an atomic prior^33^ enables Gibbs sampling and yields a sparse NMF solution, with matrix elements able to be exactly zero in cases where 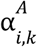 and 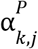 are identically zero. For our purposes, we consider convergence to be attained when additional iterations no longer reduce the chi-squared value, that is, when it has stabilized. Previously, we have found robust performance on scRNA-seq for α=0.01 and convergence after approximately 50,000 iterations for both equilibration and sampling^12^. Therefore, we use this algorithm and these parameters for the examples in this protocol.

### Dimensionality estimation

The solutions learned by NMF depend critically upon the dimensionality *k* of the factorization, which is equal to the number of patterns, and therefore also equal to the number of columns in the amplitude matrix and the number of rows in the pattern matrix. How to estimate the optimal dimensionality remains an open question in the field of unsupervised learning. In performing robustness analyses to estimate *k*, we have found that these statistics may also have local minima for pattern robustness at different dimensions. In this case, greater resolution of multiple biological components often occurs at the second, higher value of *k* for which stability is first lost. Moreover, these two local optima may both reflect distinct, hierarchical information about the underlying biological system with the dataset. For example, in a bulk genomics dataset of head and neck tumors we found that NMF at *k* = 2 separated tumor and normal samples whereas NMF at *k* = 5 separated known head and neck cancer subtypes^47^. Therefore, it may be that there is no one true *k* for NMF, but rather different biological differences are uncovered at different dimensions. Similar observations have been found in genomics analysis with other unsupervised learning techniques, including recently with autoencoders^48^.

Based upon these findings, we recommend and describe dimensionality estimation based upon tested that require solving for a range of *k* values. Linking solutions from multiple dimensionalities based on similarity and gene membership can not only provide information about robustness, but also uncover hierarchical relationships between patterns^3,19,46,49^. Additionally, the cophenetic correlation coefficient can be used to assess the stability of sample clustering at a given dimensionality as described in Brunet *et al*^1,50^. When the clustering within a dimensionality is perfectly stable the cophenetic correlation coefficient equals 1. Thus, increasing the dimensionality until the magnitude of the cophenetic correlation coefficient is >1 can determine the maximum *k* at which cluster stability is preserved.

For the workflows and data sets we present here, we chose nPatterns = 8 based on multiple runs at a range of nPatterns from 8 to 12. We settled on 8 patterns as marker gene analysis of the results using nPatterns = 10 and nPatterns = 12 showed that patterns learned at the higher dimensionalities also represented the same biological processes in the nPatterns = 8 results based on overrepresentation analysis of pattern marker genes with hallmark gene sets while additional patterns were learned. Thus the 8 pattern results were chosen for further analysis as they captured processes that also predominated at higher dimensionality while not diluting signal across a larger number of patterns.

We note that regardless, *k* must be far less than either dimension of the input dataset to yield theoretically identifiable solutions from NMF. However, similarly to other machine learning paradigms, the stability of solutions beyond this theoretical upper bound has been observed. Thus, is it likely that NMF may also experience a double-descent phenomena.

### Analysis and visualization of inferred cellular features in the pattern matrix

Single-cell experiments can provide measurements associated with numerous features of biological systems, including cell type, cell state, temporal transitions, cell cycle and metabolic states, and spatial localization^51^. Yet, the data also includes numerous technical artifacts from features, notably batch effects between libraries, dissociation protocols, and dropout^52,53^. A critical advantage of NMF for scRNA-seq data is its ability to learn separate patterns associated with each of the biological and technical features from a single analysis^3^. Nonetheless, uncovering these features from an NMF analysis of scRNA-seq data depends critically upon relating the weights of the matrix elements for each row of the pattern matrix and amplitude matrix to the biological feature or technical artifact that they represent^24^.

The most direct means of assessing the biological significance of each pattern is to correlate its values with annotations of the experimental conditions or cell type calls in the single-cell data. However, these statistics will not delineate the cellular heterogeneity within these conditions that incentivize the use of single cell data in these studies. Therefore, visualization is a critical component of this biological interpretation of the pattern matrix (Fig 3). Dimensionality reduction tools such as t-SNE or UMAP are used for visualizing single-cell analysis, and in the case of CoGAPS, they can be used for interpreting patterns in low-dimensional space. Dynamic transitions are then apparent from high pattern weights in intermediate states between cell types or areas of high RNA velocity^54^. These dynamics will also often be apparent along pseudotime trajectories. Thus, correlation or linear models associating pattern weights to pseudotime trajectories can be used to quantify these relationships. This is an especially powerful way to support the supervised pseudotime trajectories with findings from data driven unsupervised NMF techniques. A critical advantage of NMF is its ability to learn the interrelationships between cell type and experimental conditions which are not readily apparent from the visualizations used in a typical single-cell analysis workflow. Statistical tests such as MANOVAs, t-tests, or other factor based test of the pattern weights for these conditions with the experimental covariates such as treatment, condition, age, sex, etc. can assess the significance of these learned relationships.

**Figure 3.**
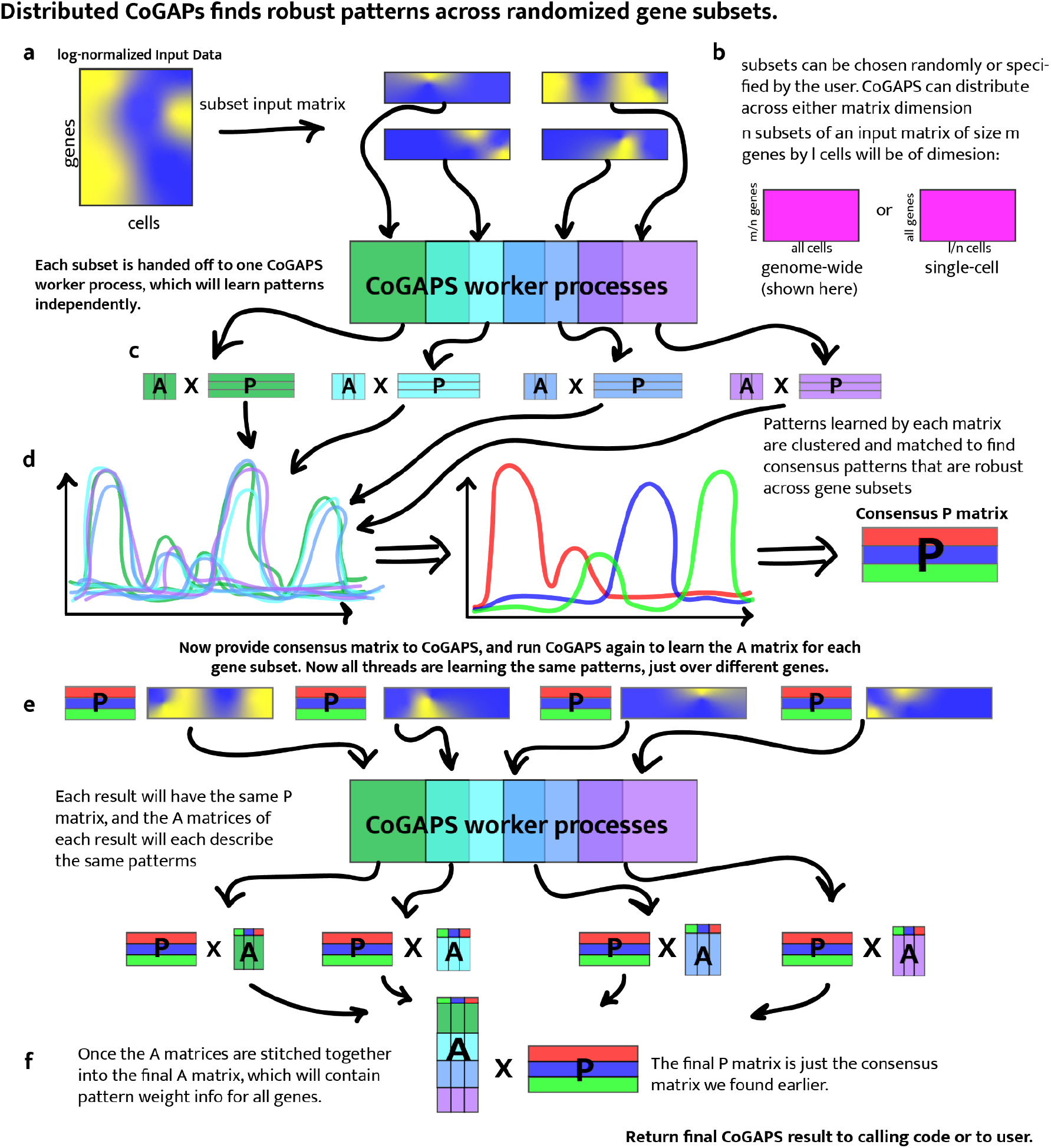
Distributed CoGAPS finds robust patterns across randomized gene or sample subsets. (a) Subsetting is performed to break the input matrices into smaller components that can each be handed off to a child process for NMF. (b) Subsetting for parallelization can be performed across either matrix dimension. (c) Each data subset yields its own NMF result. (d) To identify the patterns that manifest themselves consistently across all NMF results, clustering is performed and a consensus matrix is generated. (e) NMF is now run again on the same data subsets, this time with the consensus matrix provided as a ground truth from which the other matrix can be learned. This run is significantly faster than the first. (f) Once,Now that all threads have been forced to learn the same patterns, the portion of the NMF result that was not fixed can be stitched together to yield the final solution.

**Figure 4.**
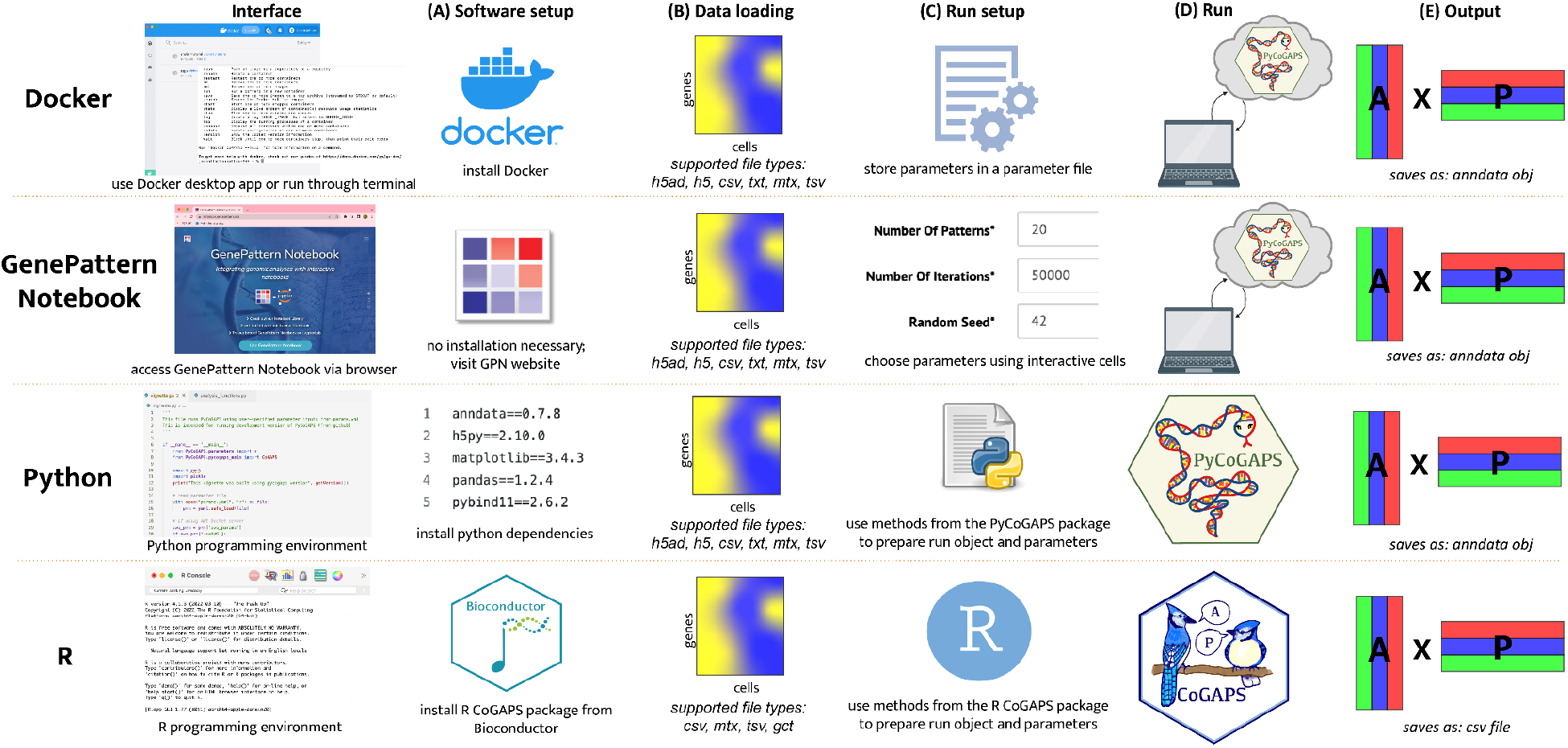
Comparison and Overview of PyCoGAPS/CoGAPS workflows.

**Figure 5.**
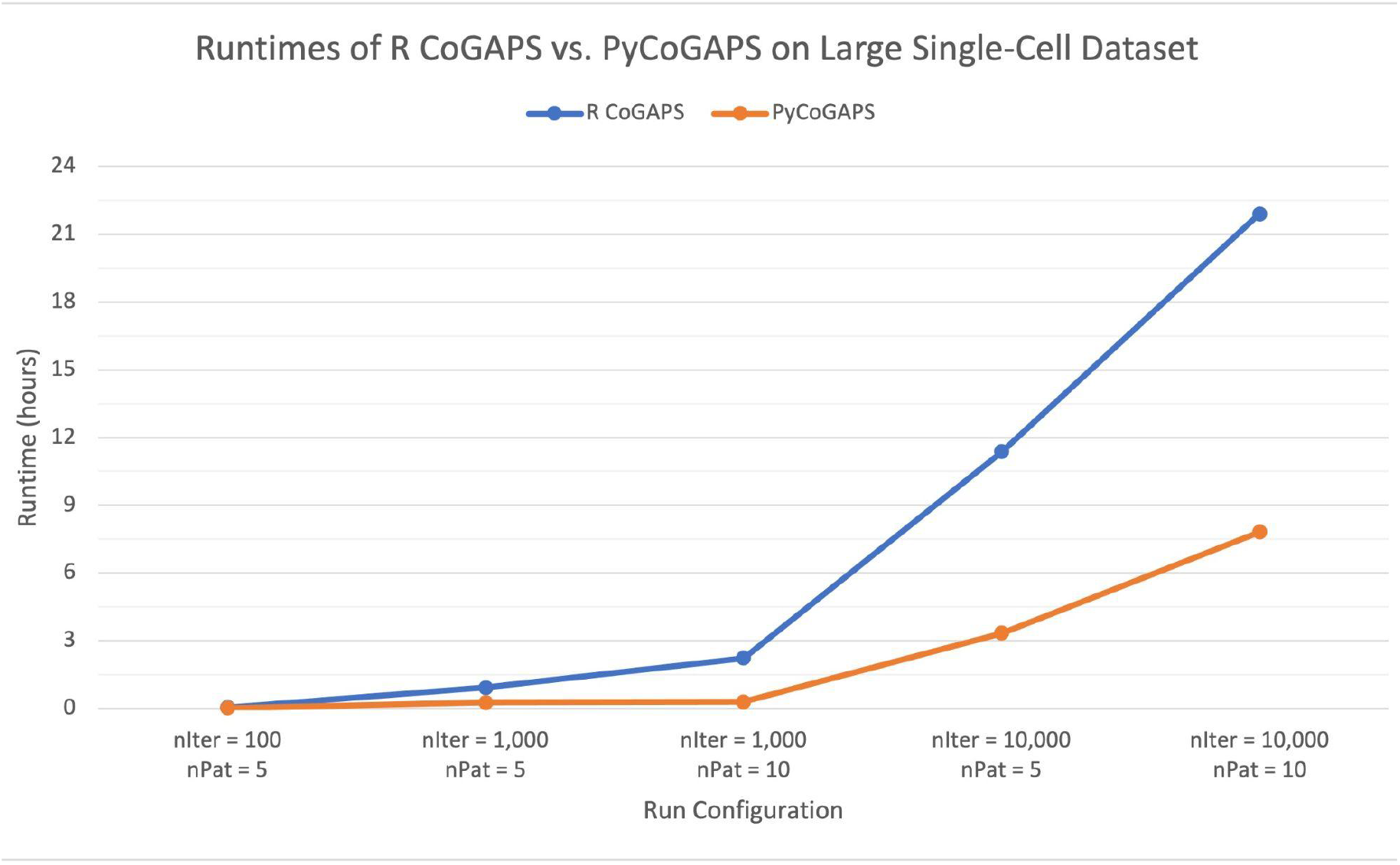
**Comparison of runtimes of R CoGAPS versus PyCoGAPS on GSE98638 data. PyCoGAPS is faster than R CoGAPS, and significantly outperforms R CoGAPS as the number of iterations and patterns increases**.

### Assessing the biological function of gene signatures from the amplitude matrix

Association of genes and pathways with the features learned from NMF analysis facilitates annotation to inform biological interpretation and hypothesis generation. For each row of the pattern matrix, there is a corresponding column in the amplitude matrix containing gene weights for the learned feature that can be used for these associations^1^. Each feature can be associated with biological processes or pathways by performing gene set analyses of the gene weights in each column of the amplitude matrix with pre-annotated sets (Fig 3) such as those curated in MSigDB^55^. In the case of Bayesian methods such as CoGAPS, these set statistics have been developed to leverage Z-scores that account for the posterior distribution of the amplitude matrix^2,19^.

An advantage of NMF for pathway discovery is its ability to highly weigh one gene in multiple columns of the amplitude matrix, reflecting the natural promiscuity and multipurpose nature of so many genes that are active in multiple biological processes, pathways, or cell types. However, this tends to hinder the identification of unique genes associated with each of the learned patterns. These marker genes are essential to define biomarkers of the learned process and prioritize candidates for experimental validation. Statistics that instead quantify the unique association of genes with each column in the amplitude matrix can be used for this analysis^6^. For example, the patternMarker statistic in CoGAPS ranks genes according to this unique association by ranking every gene for every pattern by pattern amplitude, and then iteratively matching genes to their most relevant pattern, returning a list of “marker genes” for each pattern which can then be used to interpret their biological significance. We note that often NMF analyses yield one ‘flat’ pattern that is roughly constant across all cells, accounting generally for highly expressed genes^56^. This pattern, while useful in other ways, should be excluded from the calculation of the patternMarker statistic to avoid falsely thresholding highly expressed genes. Creating a heatmap of the input data with genes ordered by their rank for each pattern can provide a clear visualization of the learned patterns^56^.

### Finding robust patterns using consensus across parallel sets

One limitation to the Bayesian structure of CoGAPS over other NMF approaches is the computational costs of numerous iterations to estimate the distribution of the amplitude and pattern matrices. The computational cost of these iterations increases as a function of the size of the dataset. To overcome this computational cost, CoGAPS supports a “distributed” mode of running in which the input data is sampled into *n* subsets of genes across every cell (genome-wide mode) or *n* subsets of cells across every gene (single-cell mode) in an embarrassingly parallel manner^25,46^. Subsetting can be performed randomly, explicitly, or using weighted assignments to ensure an even distribution of cell types among sample subsets. These supervised options are critical for users who wish to discover patterns associated with a rare cell type. For example, a pattern representing semistable cell state transitions from normal to cancer was identified in the PDAC data by enriching for epithelial cells.

Next, CoGAPS is run on each input matrix and these results are clustered and transformed into a smaller set of consensus patterns, the rationale being that robust biological patterns will manifest themselves across multiple subsets of genes or cells. For randomly sampled independent subsets, the robustness of the learned patterns can be statistically quantified. The resulting consensus matrix (either A or P depending on the mode) is then given as input to another CoGAPS run across the same subsets. This forces each thread to learn only the non-fixed matrix, so the patterns returned from this run will all be directly comparable across subsets (ie, pattern 1 in subset 1 is the same as pattern 1 in subsets 2, 3, and 4). This process enables the results to be combined into complete A and P matrices that factor the original input matrix. By using this consensus process, not only is there a significant increase in computational efficiency, but also an increased robustness of the final solution^3,46,57^.

### Multiomic methods

Coupled NMF methods^47,58^ that simultaneously decompose multiple datasets can reveal shared features with the visualizations and *post hoc* statistics on the output matrices as described above. While applicable for multi-omics analysis^58^, the implicit assumptions of these coupled methods may not accurately model timing differences between datasets or features unique to one. As an alternative, transfer learning methods that project the gene weights from the amplitude matrix learned in one source dataset onto the other datasets to compare the use of features in this new dataset. We have found that only biological features, not technical, successfully transfer between related datasets and enable comparison between data platforms, species, tissues, and molecular modalities^3,506^. This transfer learning approach can be used to annotate features in the original input source dataset based on information from the new target dataset. For example, our NMF analysis of scRNA-seq data from epithelial cell state transitions resulting from fibroblast interactions were preserved in co-culture scRNA-seq data from an *in vitro* organoid model. In the context of cancer immunotherapy, this approach also enables the discovery of preserved cell state transitions from therapy that are shared between preclinical models and human tumors^59^. Likewise, this transfer learning approach can enable integration with spatial single-cell data or imaging spatial to enable mapping of non-spatially resolved single-cell data^15,60^.

### Limitations

While we focus on NMF analysis with CoGAPS in this protocol, we note that many of the visualization and interpretation steps are also applicable to results obtained with alternative factorization methods and that there is no universal consensus as to the most robust factorization method for single-cell data.

The unsupervised nature of NMF can limit the interpretation of features to prior knowledge or annotations of the biological system measured with the single cell data. New techniques for independent assessment of biological robustness and interpretation are essential for biological discovery. Although beyond the scope of this protocol, NMF analyses comparing multiple datasets enable assessment of the robustness of learned features and discover new relationships between distinct biological contexts^10^.

## User guides: Three vignettes to run CoGAPS using Python, R, and GenePattern Notebooks

The following section contains three vignettes for NMF analysis with CoGAPS across distinct software platforms. The first demonstrates how to run the new PyCoGAPS, first by directly using the package and then via GenePattern Notebook and Docker. Finally, we provide an updated vignette workflow for our existing R interface, tailored toward the single-cell use case. All of these frameworks are functionally equivalent and share the same backend, so the user’s choice of interface should depend on factors such as computing performance and familiarity with the programming language.

### Data

Each of the following workflows will demonstrate how to run NMF (CoGAPS) on a publicly-available single-cell RNA seq dataset profiling thousands of epithelial cells in the pancreatic cancer and control pancreas samples, as well as how to understand, interpret, and use the patterns learned by CoGAPS^26^.

The data can be downloaded here: https://ngdc.cncb.ac.cn/gsa/browse/CRA001160

### Hardware

CoGAPS is usable on most laptops, personal computers, and computing clusters. We recommend first performing a coarse grained parameter sweep before fine tuning to maximize biological discovery and minimize the computational load.

### Software

Operating system: MacOS, Linux, Windows, or the Ubuntu subsystem for windows, which can be configured according to these instructions: https://docs.microsoft.com/en-us/windows/wsl/install

To use the R interface, you will need:

- R (we recommend installing it along with Rstudio, available here: https://www.rstudio.com/products/rstudio/download/
- CoGAPS, available at https://www.bioconductor.org/packages/release/bioc/html/CoGAPS.html and https://github.com/FertigLab/CoGAPS

To use PyCoGAPS, you will need:

- If using Docker option:
  - Docker, available here: https://docs.docker.com/get-docker/
- If using Python scripts option:
  - Python v. 3.8 or newer
  - PyCoGAPS, available here: https://github.com/FertigLab/pycogaps
  - A C++ compiler
    - Linux: comes standard with most if not all distributions
    - MacOS: ensure XCode is installed on your machine. If using the M1 chip, we recommend updating your software to at least MacOS Monterey 12.2.1 as it fixes a crucial issue with compiler linkages.
    - Windows: you may need to install Microsoft Build Tools. If you experience significant issues during compilation, we recommend building CoGAPS on the Ubuntu subsystem, which is available on the Windows application store.

To use the GenePattern Notebook, you will need:

- A web browser

**Table 1.**
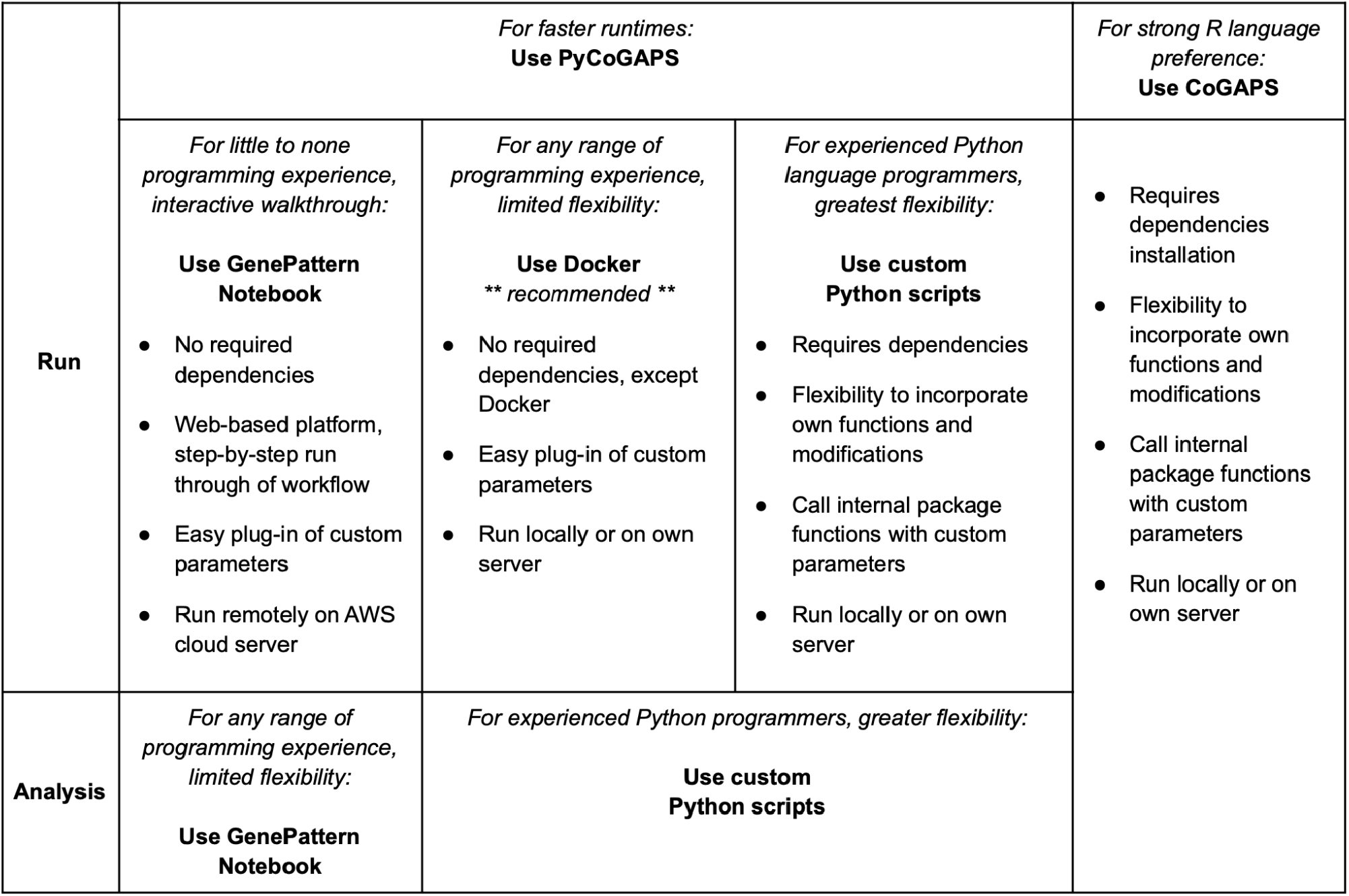
Decision table for selecting which PyCoGAPS/CoGAPS workflow to implement.

## PyCoGAPS, the Python interface to CoGAPS

This vignette aims to demonstrate the basic usage and functionality of the PyCoGAPS package, a new Python interface to the CoGAPS algorithm. PyCoGAPS was developed to make CoGAPS more accessible to researchers who rely primarily on Python and to speed up processing time by making use of Python’s innately more efficient and lightweight handling of memory and large objects. It duplicates the functionality found in the R interface, with additional visualization options and a few minor modifications necessitated by technical differences between the two languages.

PyCoGAPS can be used in a Python script like any other package, or automatically deployed and run locally or remotely using Docker. The following two vignettes will demonstrate first how to set up a CoGAPS run and analyze the result in a typical script, and the second is a comprehensive guide to using our Docker image. Setup guide and documentation can also be found in the GitHub readme: https://github.com/FertigLab/pycogaps/tree/master

### Part one: Using the PyCoGAPS Python package

#### Installation

To install, please clone our GitHub repository as follows:

~~~
git clone https://github.com/FertigLab/pycogaps.git --recursive
cd pycogaps
python setup.py install
~~~

When PyCoGAPS has installed and built correctly, you should see this message:

~~~
Finished processing dependencies **for** pycogaps==0.0.1
~~~

Which means PyCoGAPS is ready to use! You may need to install some Python dependencies before everything can build, so don’t be deterred if it takes a couple of tries to install.

### Configuring a PyCoGAPS run

If you are using the distributed CoGAPS option, all of your calling code must be wrapped in a check to make sure the main thread is running. Otherwise, child processes will attempt to call this code each thinking they’re the “base” thread and resulting in complete disaster. This is due to a quirk of Python’s multiprocessing library and wrapping your code in this check is the best solution we have found.

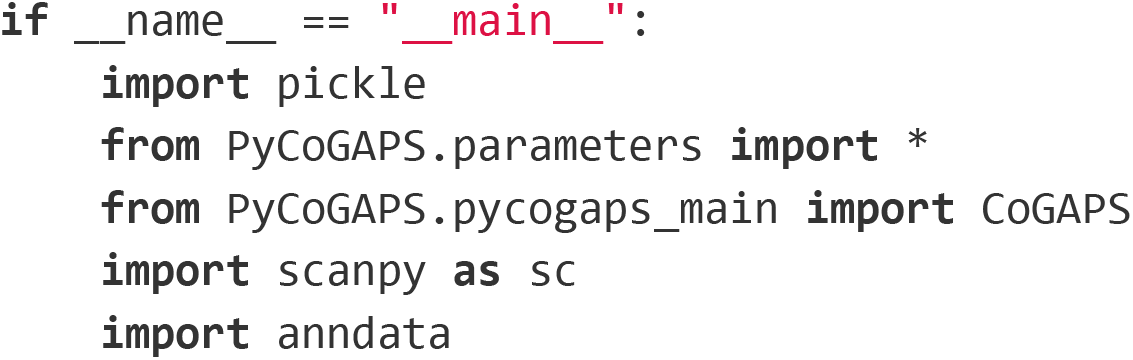

We have decided to make it possible to import different modules of PyCoGAPS separately so that nobody needs to spend their time resolving dependency issues for something they don’t need. For instance, if you don’t want to use our graphical analysis functions, then you need not import them and therefore you will not need to install scipy or seaborn.

Now we read in the anndata object containing our data. CoGAPS can handle multiple data formats, but we strongly recommend converting your data to Anndata format using the anndata package^61^ or another utility designed for translating between data structures^62^. The returned object will be in Anndata format.

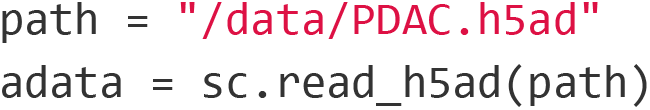

Log-normalize data (strongly recommended to normalize data before running CoGAPS). Here we make use of the scanpy package^63^, as we will continue to do throughout the Python workflow. Again, note that any transformation or scaling you choose to perform on your count matrix must result in all non-negative values due to the core constraint of NMF.

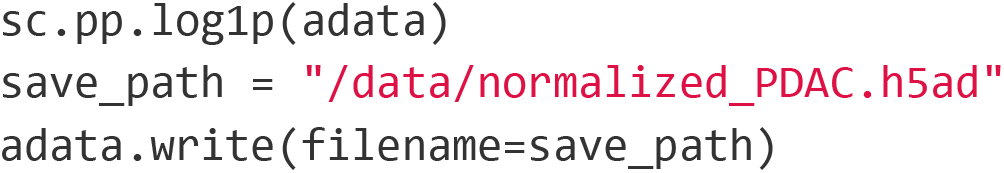

Next we create a parameters object that stores run options in a dictionary format. Note that the easiest way to decrease runtime is to run for fewer iterations, and you may want to set nIterations=1000 for a test run before starting a complete CoGAPS run on your data. There are many more run parameters that can be set depending on your goals, but for the sake of brevity, we invite the reader to explore them in our GitHub documentation.

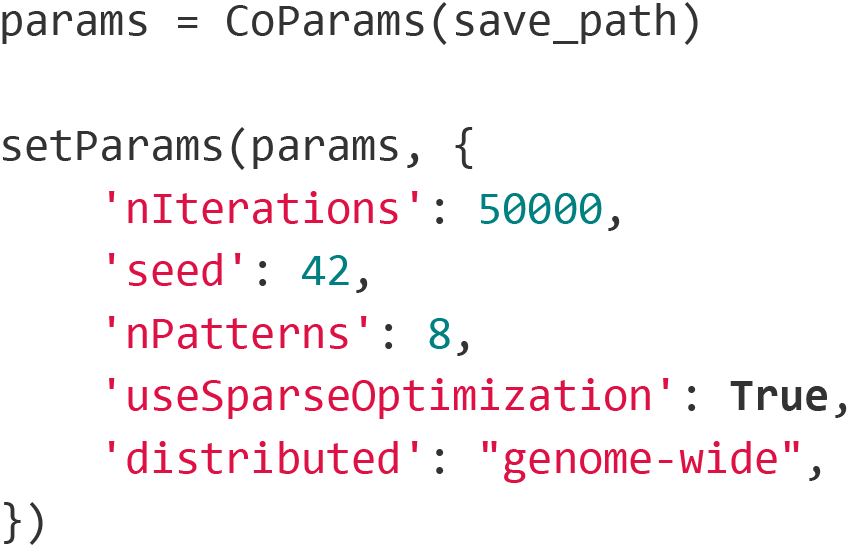

If you wish to run distributed CoGAPS, which we recommend for most cases, set the “distributed” parameter to “genome-wide” (parallelize across genes), or “single-cell” (parallelize across cells). Please see Figure 3 for a full explanation of the mechanism.

If you are running distributed, you must run this line, and you can specify how many sets will be created and parallelized across, as well as specify cutoffs for how stringently a consensus matrix is determined.

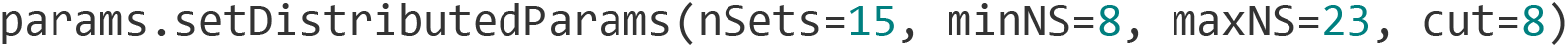

To see all parameters that have been set, call:

~~~
   params.printParams()
running genome-wide. if you wish to perform single-cell distributed cogaps,
please run setParams(params, “distributed”, “single-cell”)
setting distributed parameters - call this again if you change nPatterns
-- Standard Parameters --
nPatterns: 8
nIterations: 50000
seed: 42
sparseOptimization: True
-- Sparsity Parameters --
alpha: 0.01
maxGibbsMass: 100.0
-- Distributed Parameters --
cut: 8
nSets: 15
minNS: 8
maxNS: 23
~~~

Since CoGAPS can have quite long runtimes, we recommend timing runs and keeping rough track of how long they take to complete, in case they need to be repeated.

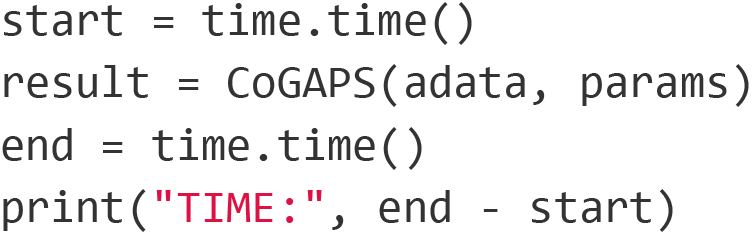

While CoGAPS is running, you will see periodic status messages saying how many iterations have been completed, the current ChiSq value, and how much time has elapsed out of the estimated total runtime.

~~~
1000 of 50000, Atoms: 5424(A), 21232(P), ChiSq: 138364000, Time: 00:03:47 /
11:13:32
1000 of 50000, Atoms: 5394(A), 20568(P), ChiSq: 133824536, Time: 00:03:46 /
11:10:34
1000 of 50000, Atoms: 5393(A), 21161(P), ChiSq: 133621048, Time: 00:03:51 /
11:25:24
1000 of 50000, Atoms: 5527(A), 22198(P), ChiSq: 137671296, Time: 00:04:00 /
11:52:06
1000 of 50000, Atoms: 5900(A), 20628(P), ChiSq: 137228688, Time: 00:03:58 /
11:46:10
~~~

When the run is finished, CoGAPS will print a message like this:

~~~
GapsResult result object with 5900 features and 20628 samples
8 patterns were learned
~~~

We strongly recommend saving your result object as soon as it returns. One option to do so is using Python’s serialization library, pickle^64^.

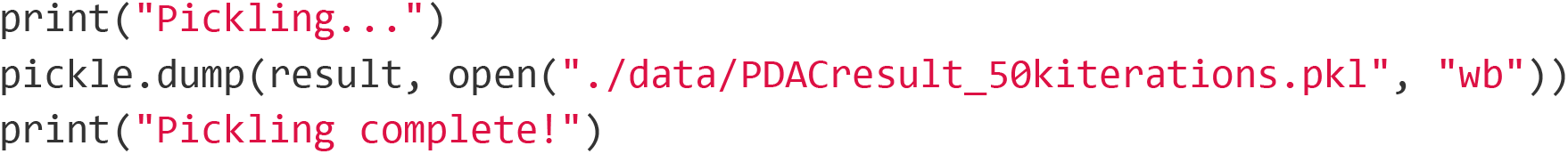

Now you have successfully generated a CoGAPS result! To continue to visualization and analysis guides, please skip to the section below titled “Breaking Down the Result Object from PyCoGAPS”

### Part two: Running PyCoGAPS using Docker

For Mac/Linux OS users, we’ll be using a Docker image, which we will pull from the Docker repository, and this contains a set of instructions to build and run PyCoGAPS. With this Docker image, there’s no need to install any dependencies, import packages, etc. as the environment is already set up and directly ready to run on your computer.

Please follow the steps below to run the PyCoGAPS vignette on Mac/Linux OS:

1. Install Docker at https://docs.docker.com/desktop/mac/install/
2. Open the Docker application or paste the following in the terminal:

~~~
docker run -d -p 80:80 docker/getting-started
~~~
3. Copy the commands and paste in terminal (Tested via Mac OX)

~~~
docker pull fertiglab/ Pycogaps
mkdir PyCoGAPS
cd PyCoGAPS
curl -O https://raw.githubusercontent.com/FertigLab/pycogaps/master/params.yaml
mkdir data
cd data
curl -O https://raw.githubusercontent.com/FertigLab/pycogaps/master/data/GIST.csv
cd ..
docker run -v $PWD:$PWD fertiglab/pycogaps $PWD/params.yaml
~~~

This produces a CoGAPS run on a simple dataset with default parameters. You should then see the following output:

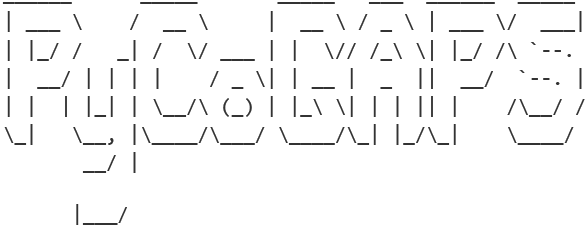

This is pycogaps version 0.0.1

Running Standard CoGAPS on GIST.csv (1363 genes and 9 samples) with parameters:

~~~
-- Standard Parameters --
nPatterns: 3
nIterations: 1000
seed: 0
sparseOptimization: False
-- Sparsity Parameters --
alpha: 0.01
maxGibbsMass: 100.0
GapsResult result object with 1363 features and 9 samples
3 patterns were learned
Pickling complete!
~~~

CoGAPS has successfully completed running and has saved the result file as `result.pkl` in a created `output/`folder.

Your working directory is the `PyCoGAPS` folder with the following structure and files:

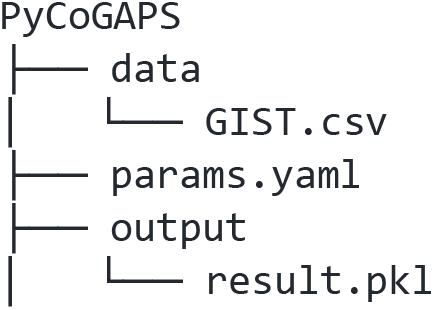

### Running PyCoGAPS on your data

Now, you’re ready to run CoGAPS for analysis on your own data with custom parameters.

In order to analyze your desired data, we’ll need to input it and modify the default parameters before running CoGAPS. All parameter values can be modified directly in the `params.yaml` file downloaded earlier.

Please see the following sections for more details on some of the parameters and options. Please follow the steps below to run PyCoGAPS with custom parameters:

1. Open `params.yaml` with any text or code editor
2. Modify the `path` parameter value by replacing the default `data/GIST.csv` with `data/your-datafile-name` Note: Make sure you have moved your data into the created `data/` folder
3. Modify any additional desired parameters and save
4. Run the following in terminal:

~~~
docker run -v $PWD:$PWD fertiglab/pycogaps $PWD/params.yaml
~~~

This step is the most time-consuming in the analysis pipeline. Timing will depend on factors such as the dimensionality of your data, the number of patterns to be learned, the number of iterations, and whether sparse optimization is enabled. For the example dataset and parameters, this step should take around 20 hours but it can vary significantly. If you change the nIterations parameter to 100, it will only take around 13 minutes, and with 1000 it will take about 80 minutes. Similar variations in timing will occur as the number of patterns is increased or decreased. The algorithm will periodically spit out status messages to let you know how far the analysis has progressed.

### Example snippet of params.yaml

The params.yaml file holds all parameters that can be inputted to PyCoGAPS. A snippet of params.yaml is shown below, where we have changed some default parameter values to our own specified example values.

~~~
## This file holds all parameters to be passed into PyCoGAPS.
## To modify default parameters, simply replace parameter values below with
user-specified values, and save file.
# RELATIVE path to data -- make sure to move your data into the created data/ folder
path: data/liver_dataset.txt
# result output file name
result_file: liver_result.pkl
standard_params:
 # number of patterns CoGAPS will learn
 nPatterns: 10
 # number of iterations for each phase of the algorithm
 nIterations: 5000
 # random number generator seed
 seed: 0
 # speeds up performance with sparse data (roughly >80% of data is zero), note this
can only be used with the default uncertainty
 useSparseOptimization: True
…
~~~

A complete list of input options and their descriptions can be found as comments in params.yaml and in Supplemental file 1, and guidelines on choosing nIterations and nPatterns is provided in the earlier portion of this manuscript.

### Multi-Threaded Parallelization

The simplest way to run CoGAPS in parallel is to modify the `nThreads` parameter in `params.yaml`. This allows the underlying algorithm to run on multiple threads and has no effect on the mathematics of the algorithm i.e. this is still standard CoGAPS. The precise number of threads to use depends on many things like hardware and data size. The best approach is to play around with different values and see how it affects the estimated time.

A snippet of `params.yaml` is shown below where `nThreads` parameter is modified.

~~~
## This file holds all parameters to be passed into PyCoGAPS.
…
run_params:
 # maximum number of threads to run on
  nThreads: 4
~~~

### Distributed PyCoGAPS

For large datasets (greater than a few thousand genes or samples) the multi-threaded parallelization isn’t enough. It is more efficient to break up the data into subsets and perform CoGAPS on each subset in parallel, stitching the results back together at the end (Stein-O’Brien et al. (2017)).

Distributed PyCoGAPS is an option that leverages the Python multiprocessing library by breaking up the data into either random or explicit sets across one dimension, running CoGAPS on each set in parallel, and then matching the patterns learned in each thread with its cognate from other runs.

In order to use these extensions, some additional parameters are required, specifically modifying the `distributed_params` in `params.yaml`. We first need to set `distributed` to be `genome-wide.` Next, `nSets` specifies the number of subsets to break the data set into. `cut`, `minNS`, and `maxNS` control the process of matching patterns across subsets and in general should not be changed from defaults. More information about these parameters can be found in the original papers.

A snippet of `params.yaml` is shown below where `distributed_params` parameters are modified.

~~~
## This file holds all parameters to be passed into PyCoGAPS.
…
distributed_params:
 # either null or genome-wide
 distributed: genome-wide
 # number of sets to break data into
 nSets: 4
 # number of branches at which to cut dendrogram used in pattern matching
 cut: null
 # minimum of individual set contributions a cluster must contain
 minNS: null
 # maximum of individual set contributions a cluster can contain
 maxNS: null
~~~

`nSets` controls how many subsets are run in parallel when using the distributed version of the algorithm. Setting `nSets` requires balancing available hardware and run time against the size of your data. In general, `nSets` should be less than or equal to the number of nodes/cores that are available. If that is true, then the more subsets you create, the faster CoGAPS will run - however, some robustness can be lost when the subsets get too small. The general rule of thumb is to set `nSets` so that each subset has between 1000 and 5000 genes or cells in order to give robust results, but ideally we would want as many cells per set as possible.

If `explicitSets` are not provided, the data will be randomly fragmented into the number of sets specified by `nSets` parameter, with the default being 4. Subsets can also be chosen randomly, but weighted according to a user-provided annotation in parameters `samplingAnnotation` and `samplingWeight`.

For Distributed PyCoGAPS, once all worker threads have started running their iterations, you will see periodic output like this:

~~~
1000 of 50000, Atoms: 5424(A), 21232(P), ChiSq: 138364000, Time: 00:03:47 / 11:13:32
1000 of 50000, Atoms: 5394(A), 20568(P), ChiSq: 133824536, Time: 00:03:46 / 11:10:34
1000 of 50000, Atoms: 5393(A), 21161(P), ChiSq: 133621048, Time: 00:03:51 / 11:25:24
1000 of 50000, Atoms: 5527(A), 22198(P), ChiSq: 137671296, Time: 00:04:00 / 11:52:06
1000 of 50000, Atoms: 5900(A), 20628(P), ChiSq: 137228688, Time: 00:03:58 / 11:46:10
~~~

### The PyCoGAPS Result Object

The CoGAPS result is returned in Anndata format. CoGAPS stores the lower-dimensional representation of the samples (P matrix) in the .var slot and the weight of the features (A matrix) in the .obs slot. If you transpose the matrix before running CoGAPS, the opposite will be true. Running single-cell is equivalent in every way to transposing the data matrix and running single-cell. The standard deviation across sample points for each matrix as well as additional metrics are stored in the .uns slots. Please refer to https://github.com/FertigLab/pycogaps#readme for complete documentation of output metrics.

### Analyzing the PyCoGAPS Result

We provide below three examples of doing a simple analysis and visualization of the PyCoGAPS result learned in the previous section. The first is directly in Python, the second uses an interactive notebook interface using the web-based GenePattern Notebook (recommended for less experienced Python/programming users), and the third uses a prewritten analysis script that can be easily edited and run directly from the command line.

### Directly using the Python package

If you wish to write a script using our package rather than using one of the preformed options, please refer to this section.

### Visualize patterns and compare with clusters and annotations using UMAP and the scanpy package

Though many bioinformaticians who use Python have likely already integrated scanpy or an equivalent package into their workflow, we include here an example of how to visualize pattern amplitudes overlaid on the UMAP plot of the cell data. It will roughly follow the steps in this tutorial: https://scanpy-tutorials.readthedocs.io/en/latest/pbmc3k.html

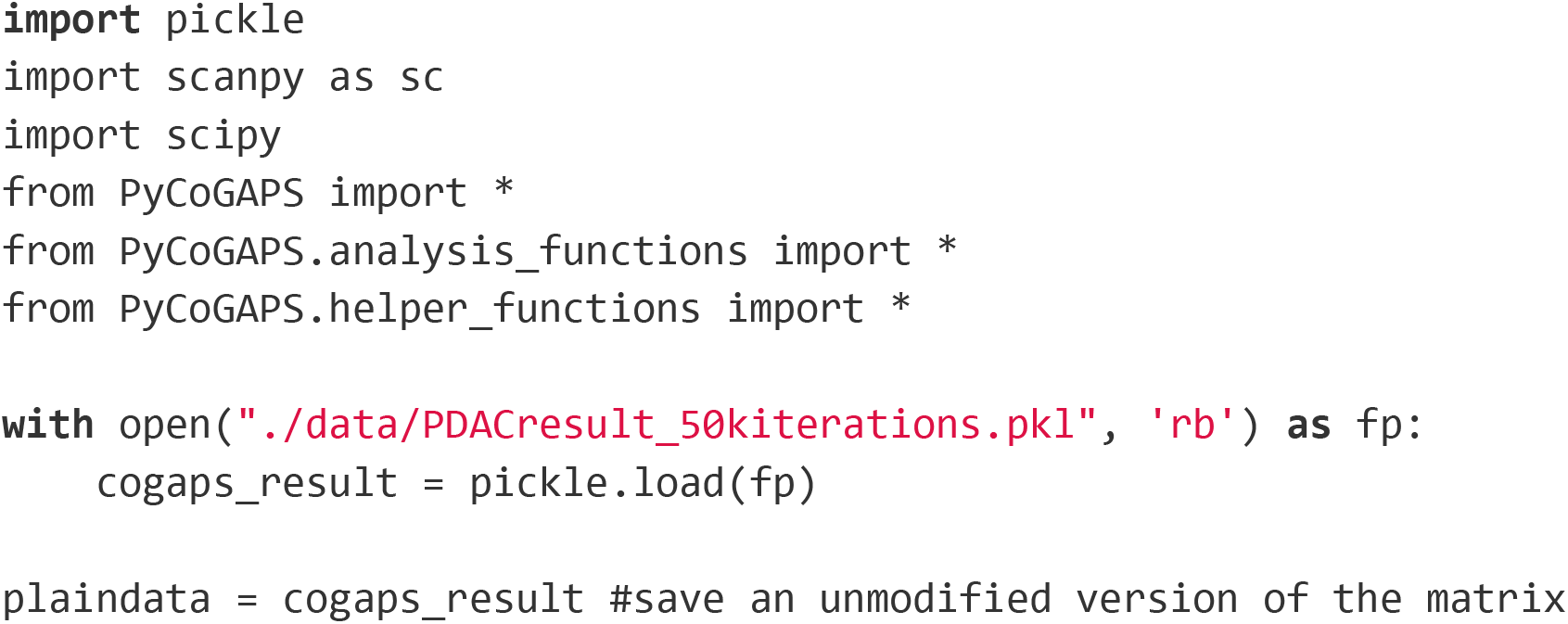

Scanpy requires genes in columns and cells in rows, so we must transpose our data object (this will also automatically transpose the obs and var matrices).

~~~
cogaps_result = cogaps_result.T
~~~

Reading in some additional annotations that I want to include in my plots later, and storing them as a column of the obs matrix (annotates each cell).

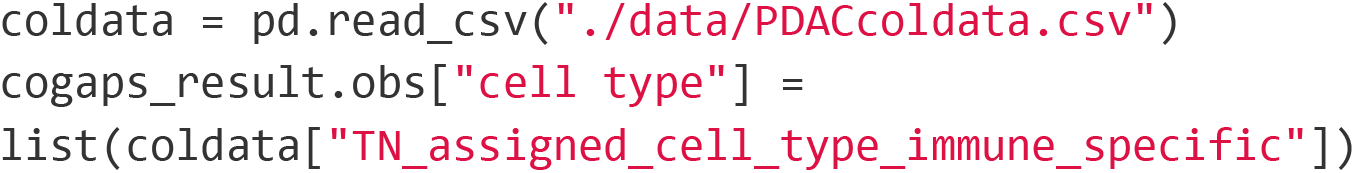

Normalize total gene count and scale data (don’t need to log normalize as we did that before running CoGAPS).

~~~
sc.pp.normalize_total(cogaps_result)
sc.pp.scale(cogaps_result)
~~~

Run PCA, learn a neighbor graph embedding, and find clusters using leiden.

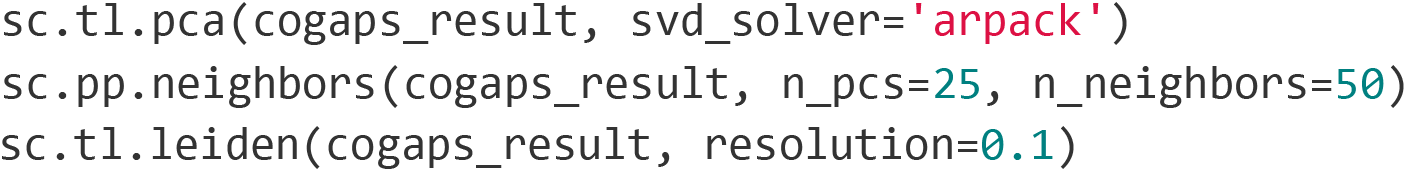

Now run UMAP and color the embedding based on pattern amplitude in each cell, along with cell identity and leiden clusters.

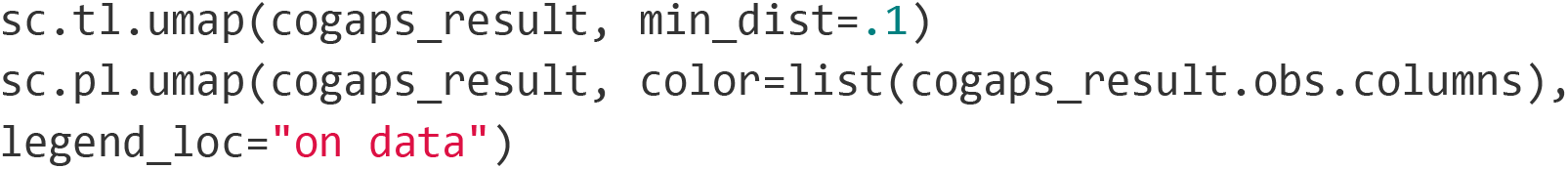

This graph is generated:

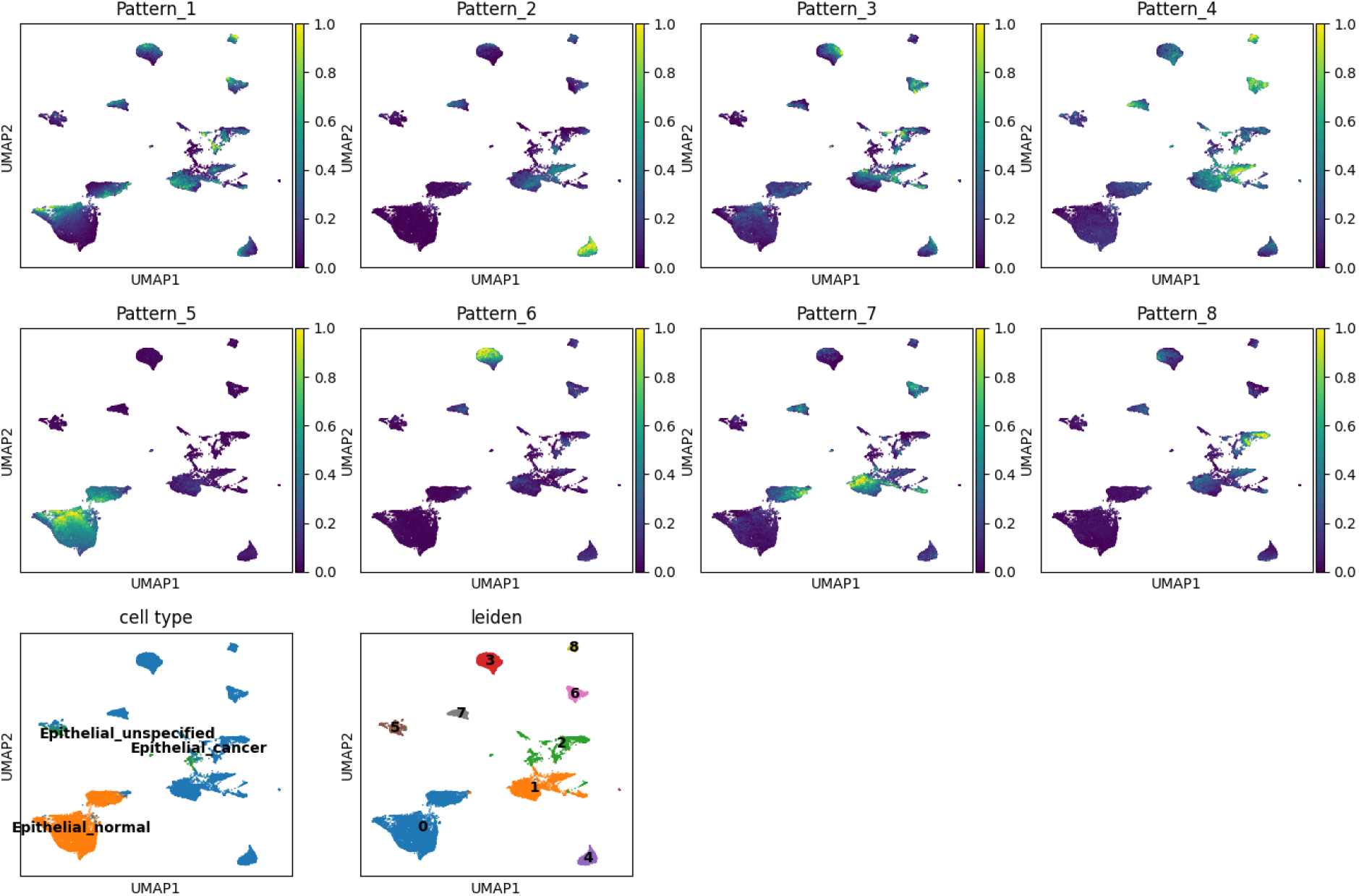

From these embeddings we can see that normal and cancer epithelial cells cluster quite separately as cluster 0, and that pattern 5 seems to describe the identity of the normal cells. There are several distinct clusters of epithelial cells from people with cancer, which are well-separated in UMAP space. Cluster 4 seems to be also described by pattern 2, and the same with cluster 3/pattern 6, cluster 2/pattern 8, clusters 7, 6, 8/pattern 4, and so forth. Pattern 7 seems to bridge clusters 0 and 1, perhaps representing a transitional pattern of gene expression. Patterns are not equivalent to clusters, but they often separate cell identity and so visually comparing them against leiden clusters can be very useful when interpreting the meaning of each pattern.

We can also visualize this using scanpy’s stacked_violin plot.

~~~
sc.pl.stacked_violin(cogaps_result, [‘Pattern_1’, ‘Pattern_2’, ‘Pattern_3’,
‘Pattern_4’, ‘Pattern_5’, ‘Pattern_6’, ‘Pattern_7’, ‘Pattern_8’],
groupby=‘leiden’)
~~~

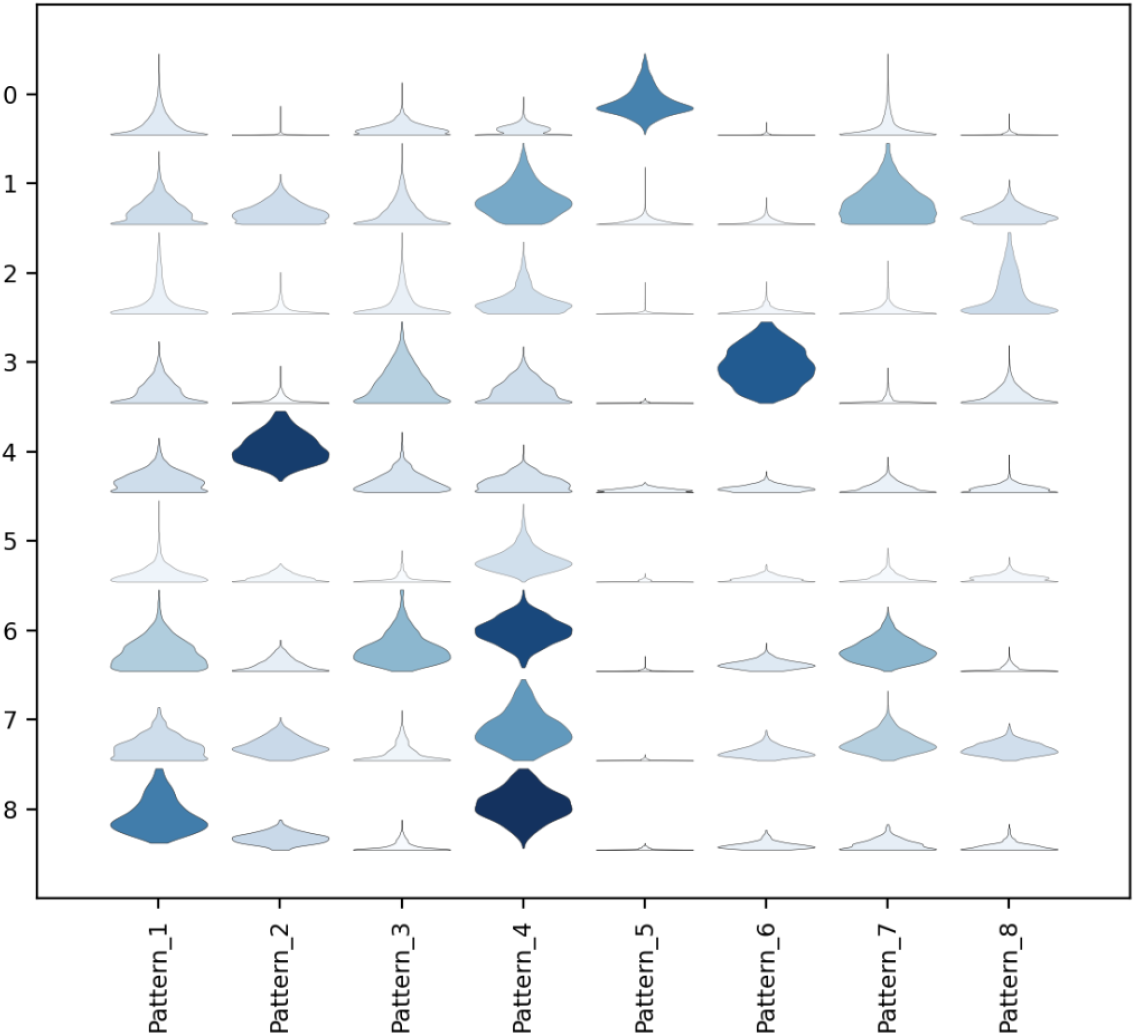

And for cancer vs. healthy cells:

~~~
sc.pl.stacked_violin(cogaps_result, [‘Pattern_1’, ‘Pattern_2’, ‘Pattern_3’,
‘Pattern_4’, ‘Pattern_5’, ‘Pattern_6’, ‘Pattern_7’, ‘Pattern_8’],
groupby=‘cell type’)
~~~

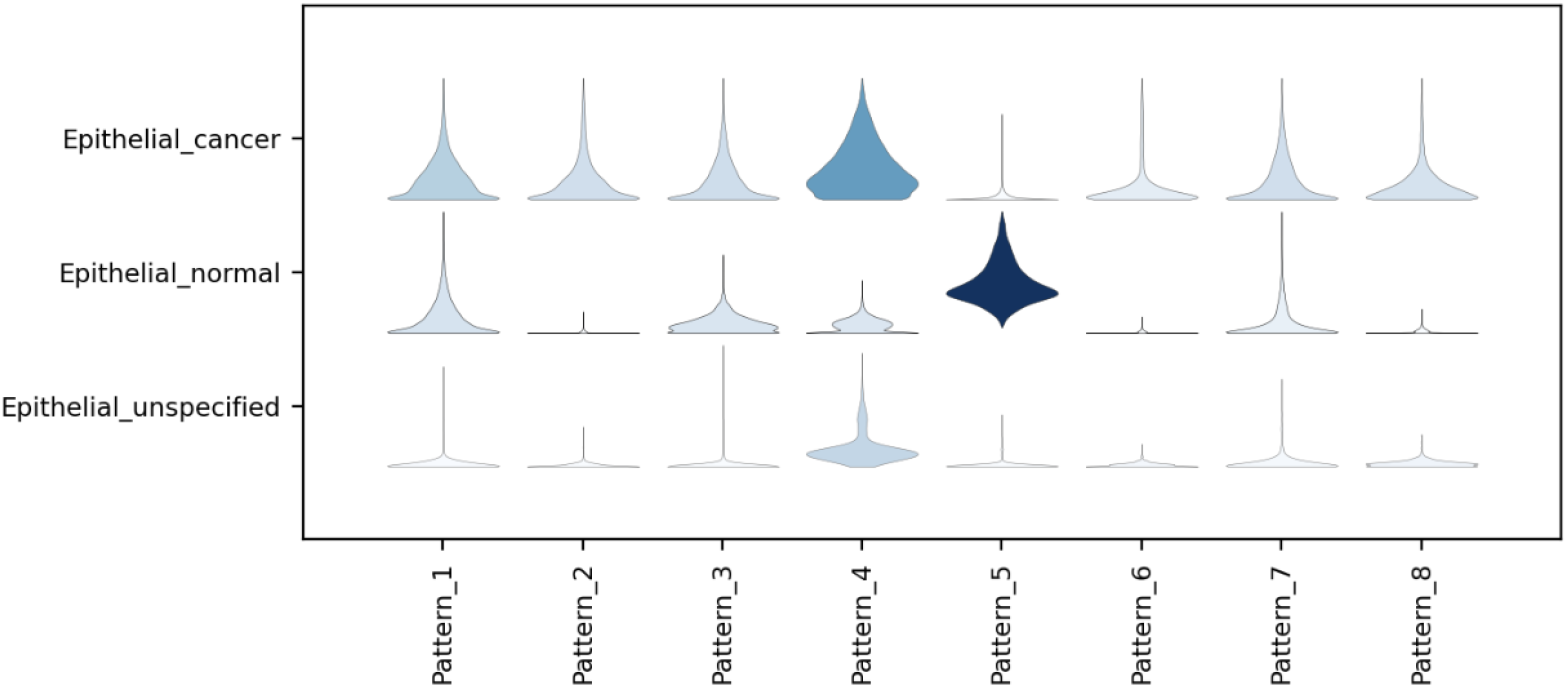

### Find markers of each pattern

Identifying genes that are strongly correlated with each learned pattern allows us to begin to decipher what biological processes or states it may represent.

The patternMarkers function, which is part of PyCoGAPS, has two modes designated by the threshold parameter. When threshold=“all”, each gene is designated as a marker of whichever pattern it is most associated with, and the number of markers will equal the number of genes (each gene is a marker of one pattern). When threshold = “cut”, a much shorter list of genes will be returned for each pattern, representing the subset of genes that are most strongly associated with that pattern. Here, we will use “cut”. Note that it may be easier to find pattern markers from the unmodified matrix we saved earlier, as patternMarkers assumes that the only columns in the obs matrix are pattern columns.

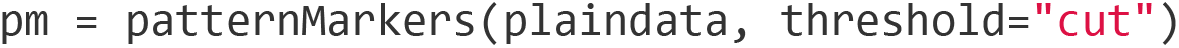

The patternMarkers function returns a dictionary with three categories: PatternMarkers, PatternMarkerRank, and PatternMarkerScores. The names are fairly self explanatory, PatternMarkers is a list of marker genes for each pattern, PatternMarkerRanks are each gene ranked by association for each pattern, and PatternMarkerScores are scores describing how strongly a gene is associated with a pattern. To view marker genes for a pattern, access it like this:

~~~
pm[“PatternMarkers”][“Pattern_1”]
[‘XRCC5’, ‘TTC14’, ‘UTRN’, ‘WNT10B’, ‘ZNF432’, ‘LEMD1’, ‘MPP7’, ‘SCARA5’,
‘KCNC4’, ‘CTTNBP2’, ‘LAMB1’, ‘USP16’, ‘PDE4DIP’, ‘SLC25A29’, ‘PSME4’,
‘CTSS’, ‘ORC5’, ‘CEP97’, ‘DEGS1’, ‘HOXA13’, ‘CTIF’, ‘C9orf16’, ‘SLC6A16’,
‘FAM135A’, ‘HECA’, ‘CAPN15’, ‘ECE2’, ‘SPATS2L’, ‘PARVG’, ‘ADAM11’,
‘SMIM13’, ‘CAMKK1’, ‘TARDBP’, ‘RFK’, ‘OSMR’, ‘RSPH9’, ‘SH2D7’, ‘ZC3H13’,
‘CAP2’, ‘CASP2’, ‘ATAD3C’, ‘CD151’, ‘LYPD5’, ‘ENTPD7’, ‘CNNM3’]
pm[“PatternMarkers”][“Pattern_7”]
[‘SPEF2’, ‘PAH’, ‘FAM117B’, ‘SFTPA2’, ‘PDLIM4’, ‘ZNF503’, ‘CITED2’,
‘GTPBP4’, ‘ZSWIM8’, ‘CHD5’, ‘TNFRSF9’, ‘CD3EAP’, ‘AMIGO2’, ‘STX3’,
‘CAMK2G’, ‘RACGAP1’, ‘SOWAHB’, ‘ABRACL’, ‘LTBP2’, ‘CDK11B’, ‘MFAP1’, ‘UNK’,
‘PLEKHH3’, ‘C1orf115’, ‘SATB1’, ‘BBOX1’, ‘SPN’, ‘UHRF1BP1’, ‘PVR’, ‘NLRP4’,
‘CAMK4’, ‘ZNF324’, ‘WWTR1’, ‘DYDC2’, ‘SHANK2’, ‘GBF1’, ‘HSPH1’, ‘VDAC2’,
‘FAM229A’, ‘COG3’, ‘RFTN1’, ‘KRT81’, ‘GLP2R’, ‘NR3C1’, ‘BNIP1’, ‘SLFN13’,
‘RABL3’, ‘TNKS’, ‘RAB30’, ‘ARHGAP21’, ‘ABTB2’, ‘ETNK1’, ‘DUS4L’, ‘PDK4’,
‘SLC35A3’, ‘ABCC5’, ‘NRK’, ‘ZNF439’, ‘TYSND1’, ‘SYAP1’, ‘GAR1’, ‘NOS3’,
‘POLR2M’, ‘SERPINI1’]
~~~

Continuing our journey, let’s collect a list of all genes ranked among the top 5 for at least one pattern.

~~~
thresh = 1
m = pm[“PatternMarkerRanks”].le(5).sum(1).ge(thresh)
top5ranked = list(pd.DataFrame(m[m==True]).index)
~~~

Now we can visualize these genes as they vary with patterns or across cell types.

~~~
sc.pl.dotplot(epiMtx, top5ranked, groupby=‘cell type’)
~~~

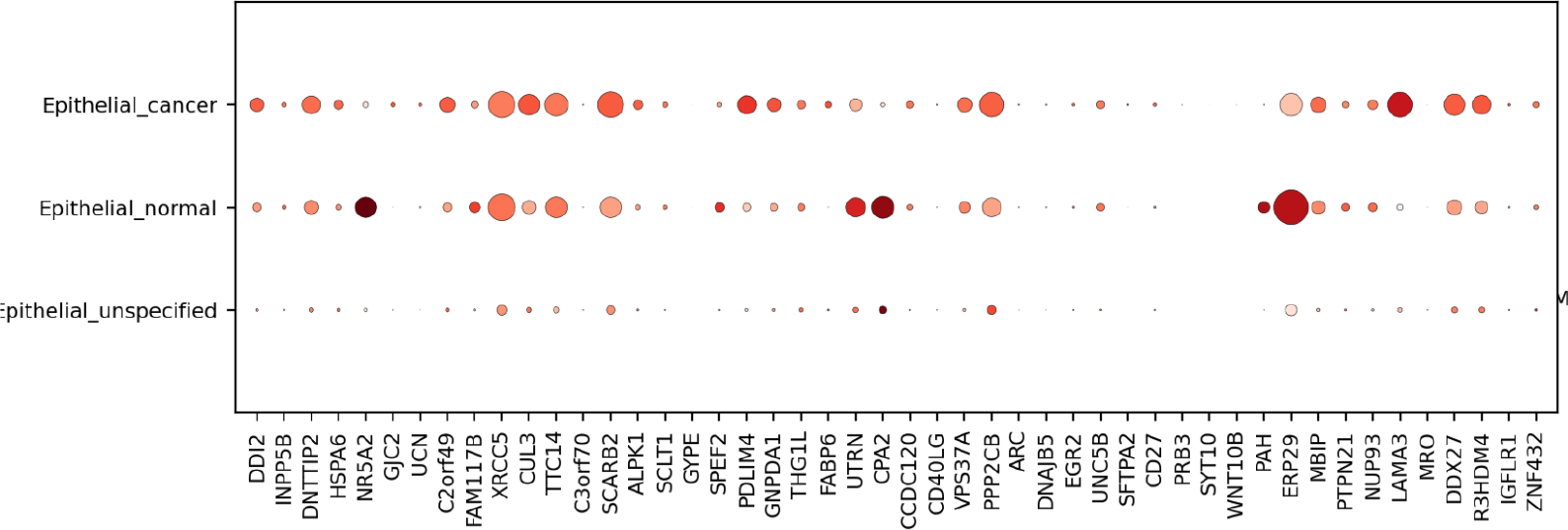

We previously noted that Pattern 7 may describe a transitional state between cancer-associated and healthy epithelial cells. We can obtain the top 15 ranked genes for pattern 7 as follows:

~~~
pattern7top50 =
list((pm[“PatternMarkerRanks”][“Pattern_7”].sort_values()[1:50]).index)
sc.pl.dotplot(epiMtx, pattern7top50, groupby=‘cell type’)
~~~

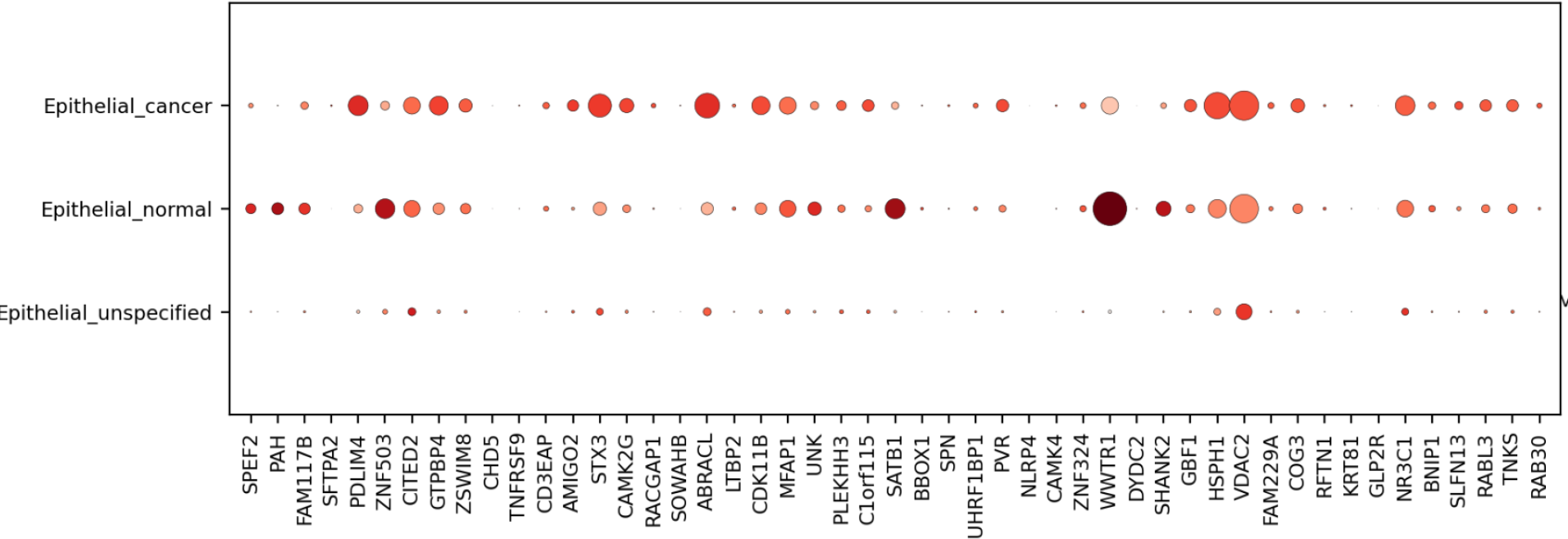

To get a cursory look at what some of these genes might be doing, I input a list of all pattern 7 marker genes into goNet^65^, an online tool that can quickly perform GO term annotation on a list of human or mouse genes (they must be entered with a newline between each gene). This is the graph that was produced:

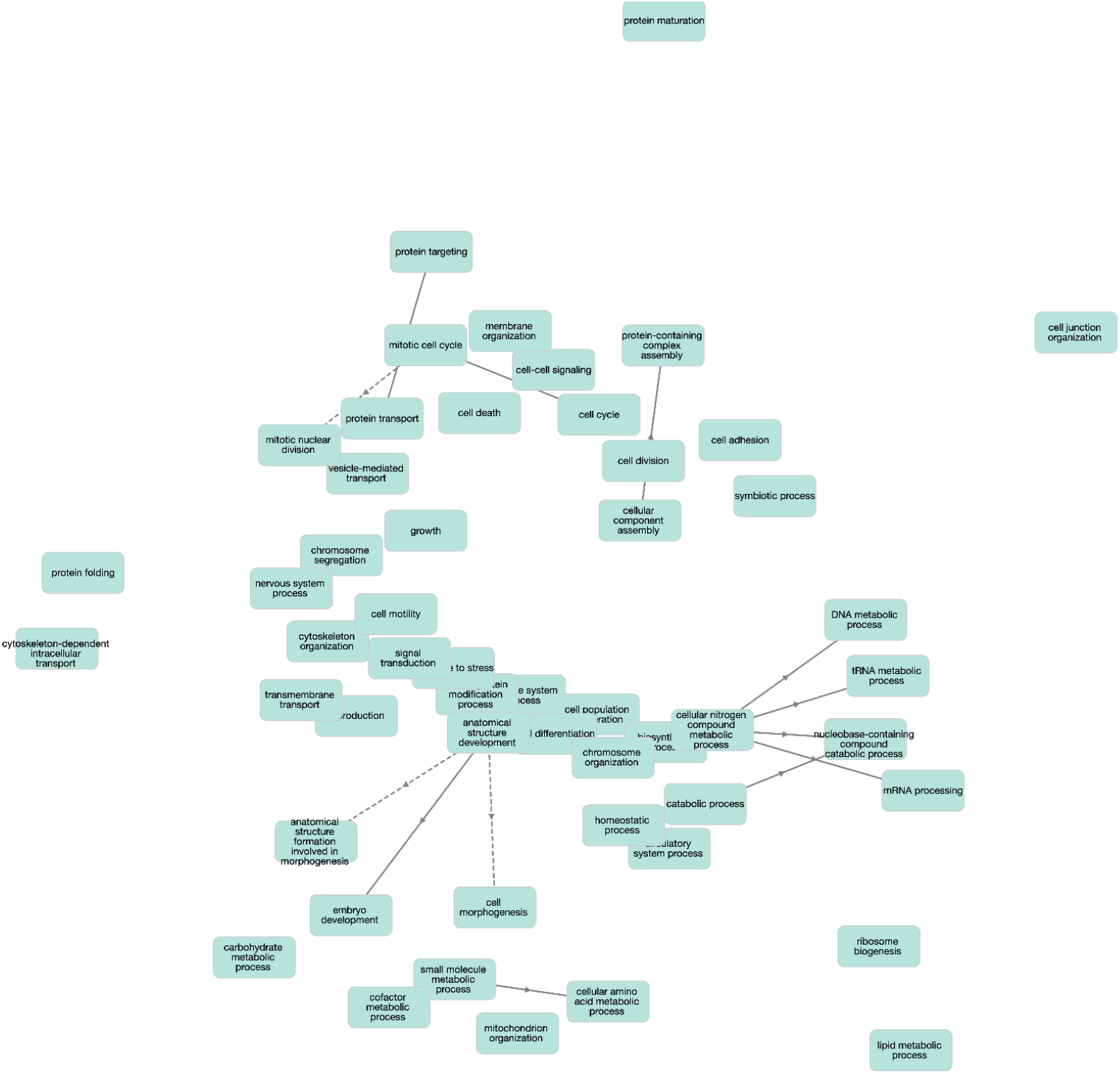

From this, we can see a striking abundance of metabolomic terms, as well as many related to proliferation and cell division. We hypothesize that pattern 7 encodes the genetic profile of a healthy epithelial cell transitioning to a more inflammatory, more proliferative, possibly more cancer-like phenotype.

### Using GenePattern Notebook

We provide an additional way of analyzing and visualizing results from your run using GenePattern Notebook, recommended for users with less programming experience.

#### Steps

1. Log in to the GenePattern Notebook workspace, http://notebook.genepattern.org. If you do not have an account, click the “Register a new GenePattern Account” button, provide the registration information, and log in. Registration for GenePattern Notebook is free.
2. Scroll to “Public Library.” You will see a list of available public notebooks.
3. In the “Search Library” box, search “PyCoGAPS.”
4. Select the “PyCoGAPS Analysis” notebook by clicking anywhere in its description and selecting “Run”. A copy of the notebook will be saved in your account.
5. Select the file titled “PyCoGAPS Analysis.ipynb” to open. The notebook describes each step in this protocol and contains cells that will allow you to execute its corresponding analyses and visualizations. Instructions for providing input datasets and setting parameters are provided in blue panels above each analysis step.
6. Follow the instructions in each blue panel, providing input data where requested.

### Python Script from Terminal

If you would like to quickly generate figures from your CoGAPS result, this Python script will run rudimentary analysis and produce visualizations. In order to analyze the data, we’ll need to make sure to install the necessary dependencies and import the built-in PyCoGAPS functions.

Make sure you’re in the `PyCoGAPS` folder, and copy the following commands in terminal, which will save plots generated from the example data in the `output/` folder:

~~~
cd output
curl -O
https://raw.githubusercontent.com/FertigLab/pycogaps/master/PyCoGAPS/analysis_functions.py
curl -O
https://raw.githubusercontent.com/FertigLab/pycogaps/master/PyCoGAPS/requirements_analysis.txt
pip install -r requirements_analysis.txt --user
python3 analysis_functions.py result.pkl
~~~

To analyze a different result, replace `result.pkl` with the path to your desired result file in the command line.

## R Interface to CoGAPS

### Requirements

CoGAPS has an interface that can be run entirely through R, with a vignette available on the page for its Bioconductor release: https://www.bioconductor.org/packages/devel/bioc/vignettes/CoGAPS/inst/doc/CoGAPS.html

### Installation

CoGAPS is a Bioconductor package and so the release version can be installed as follows:

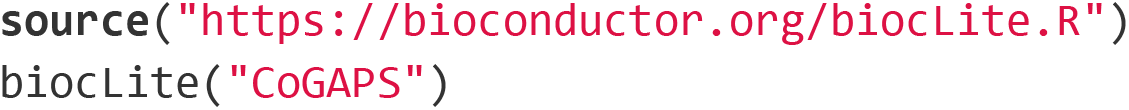

The most up-to-date version of CoGAPS can be installed directly from the FertigLab Github Repository:

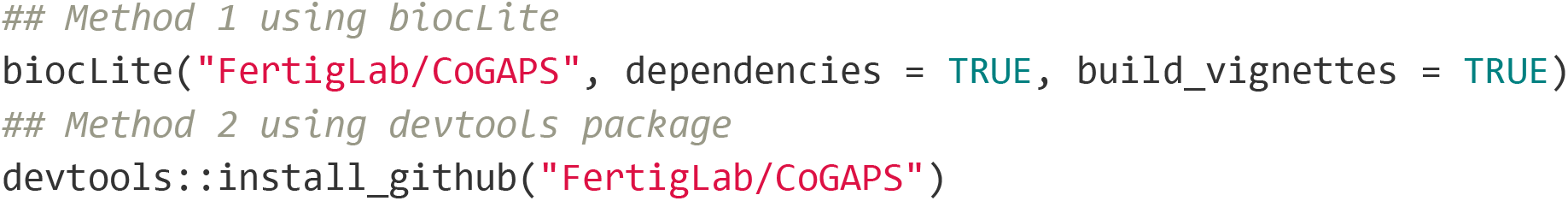

There is also an option to install the development version of CoGAPS, while this version has the latest experimental features, it is not guaranteed to be stable.

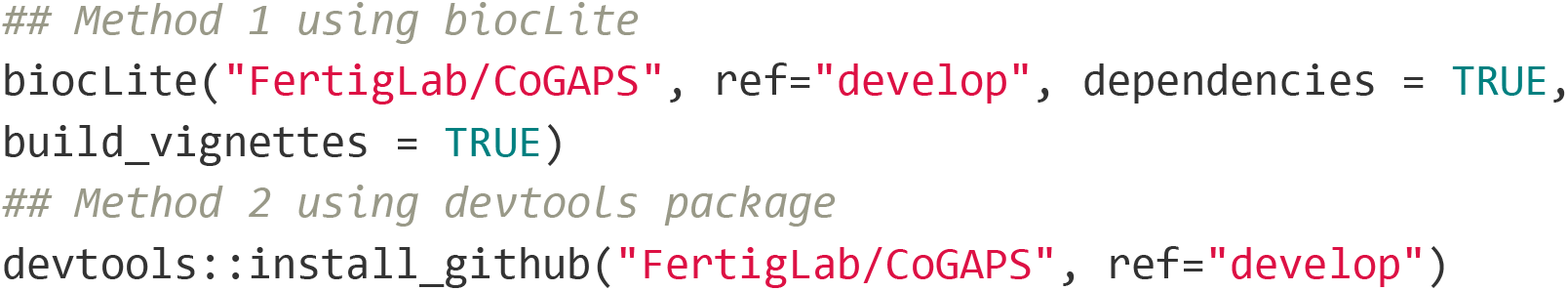

### Setting up a CoGAPS run

Import necessary libraries

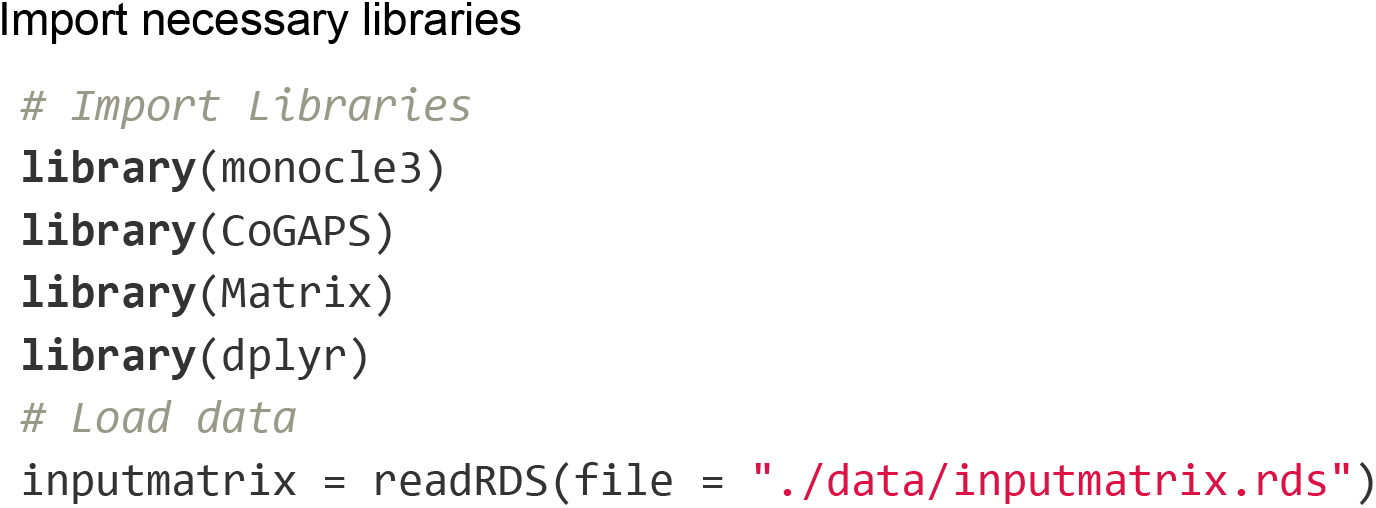

Most of the time we’ll want to set some parameters before running CoGAPS. Parameters are managed with a CogapsParams object. This object will store all parameters needed to run CoGAPS and provides a simple interface for viewing and setting the parameter values.

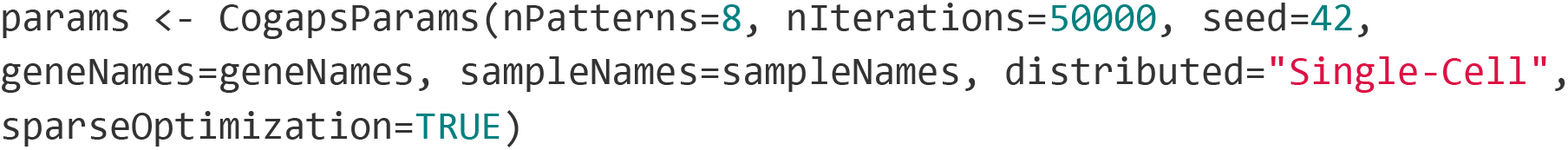

If you wish to run distributed CoGAPS, which is strongly recommended for most large datasets, you must also call the setDistributedParams function.

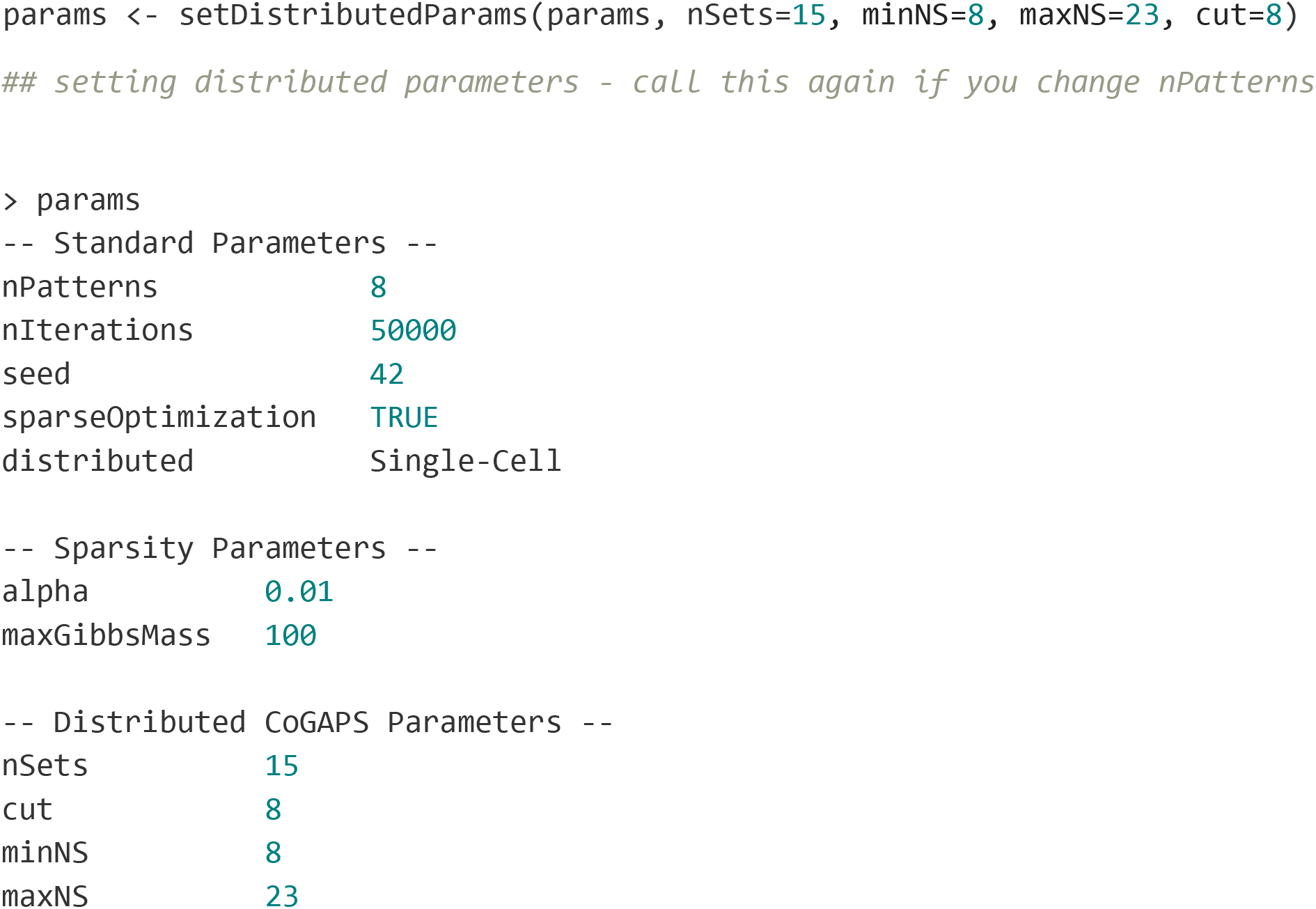

Now to start your CoGAPS run simply call:

~~~
cogaps_result <- CoGAPS(inputmatrix, params)
~~~

While CoGAPS is running it periodically prints status messages. For example:

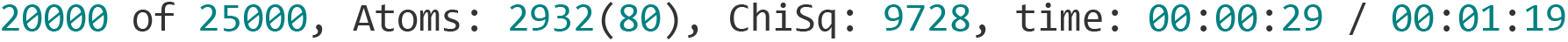

This message tells us that CoGAPS is at iteration 20000 out of 25000 for this phase and that 29 seconds out of an estimated 1 minute 19 seconds have passed. It also tells us the size of the atomic domain which is a core component of the algorithm but can be ignored for now. Finally, the ChiSq value tells us how closely the A and P matrices reconstruct the original data. In general, we want this value to go down - but it is not a perfect measurement of how well CoGAPS is finding the biological processes contained in the data. CoGAPS also prints a message indicating which phase is currently happening. There are two phases to the algorithm - Equilibration and Sampling.

### Breaking down and analyzing the CoGAPS result object

Now that the CoGAPS run is complete, it’s time to investigate the patterns it learned. It is generally a good idea to immediately save your CoGAPS result object to a file, then read it when you’re ready to work with it.

~~~
cogaps <- readRDS(“CoGAPS_output/cogapsresult.rds”)
> cogaps
[1] “CogapsResult object with 15176 features and 25442 samples”
[1] “8 patterns were learned”
~~~

The CoGAPS result object consists of:

- A and P matrices learned by CoGAPS. In this package, the A matrix of sample weights is called “**sampleFactors**” and the P matrix of gene weights is called “**featureLoadings**”.
- Standard deviation matrices **factorStdDev** and **loadingStdDev** corresponding to sampleFactors and featureLoadings
- Metadata, which contains information for the run such as how it was parallelized (**subsets**), the mean ChiSq value during the run (**meanChiSq**), and the parameters used in the run (**params**). Since the run parameters are attached to the result object, it makes it easy to keep track of the provenance of your CoGAPS results. Other information may be present in the metadata depending on your run options.

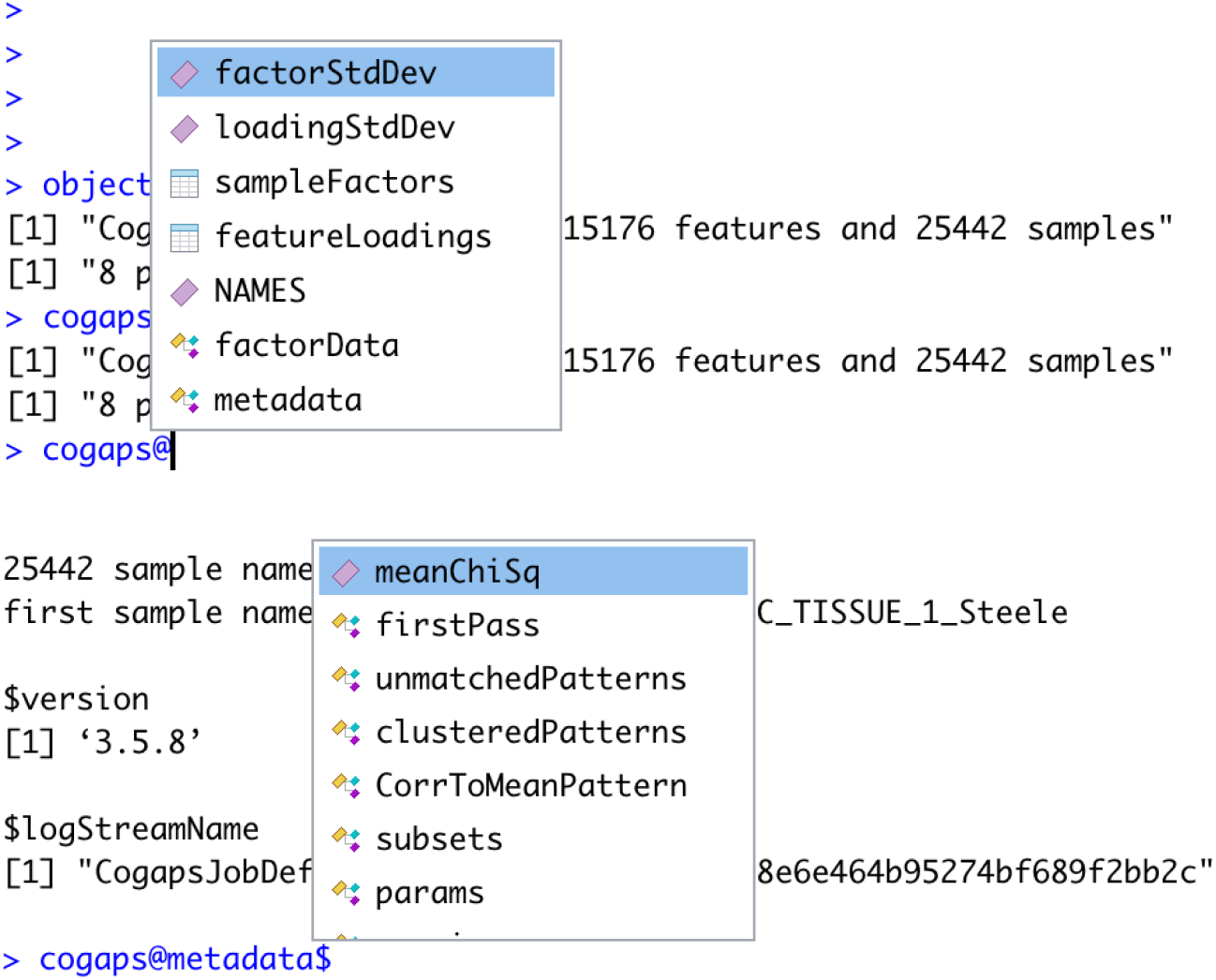

### Visualize learned patterns on a UMAP embedding

Often, it is a good idea to immediately visualize pattern weights on a UMAP because you will immediately see whether they are showing strong signal and make common sense.

Since pattern weights are all continuous and nonnegative, they can be used to color a UMAP in the same way as one would color by gene expression. The sampleFactors matrix is essentially just *nPatterns* different annotations for each cell, and featureLoadings is likewise just *nPatterns* annotations for each gene. This makes it very simple to incorporate pattern data into any data structure and workflow.

We have added a built-in function, plotPatternUMAP, which renders a UMAP embedding of your cells colored by the intensity of each pattern.

~~~
cds <- reduce_dimension(cds)
pl <- plotPatternUMAP(object, cds)
pl
~~~

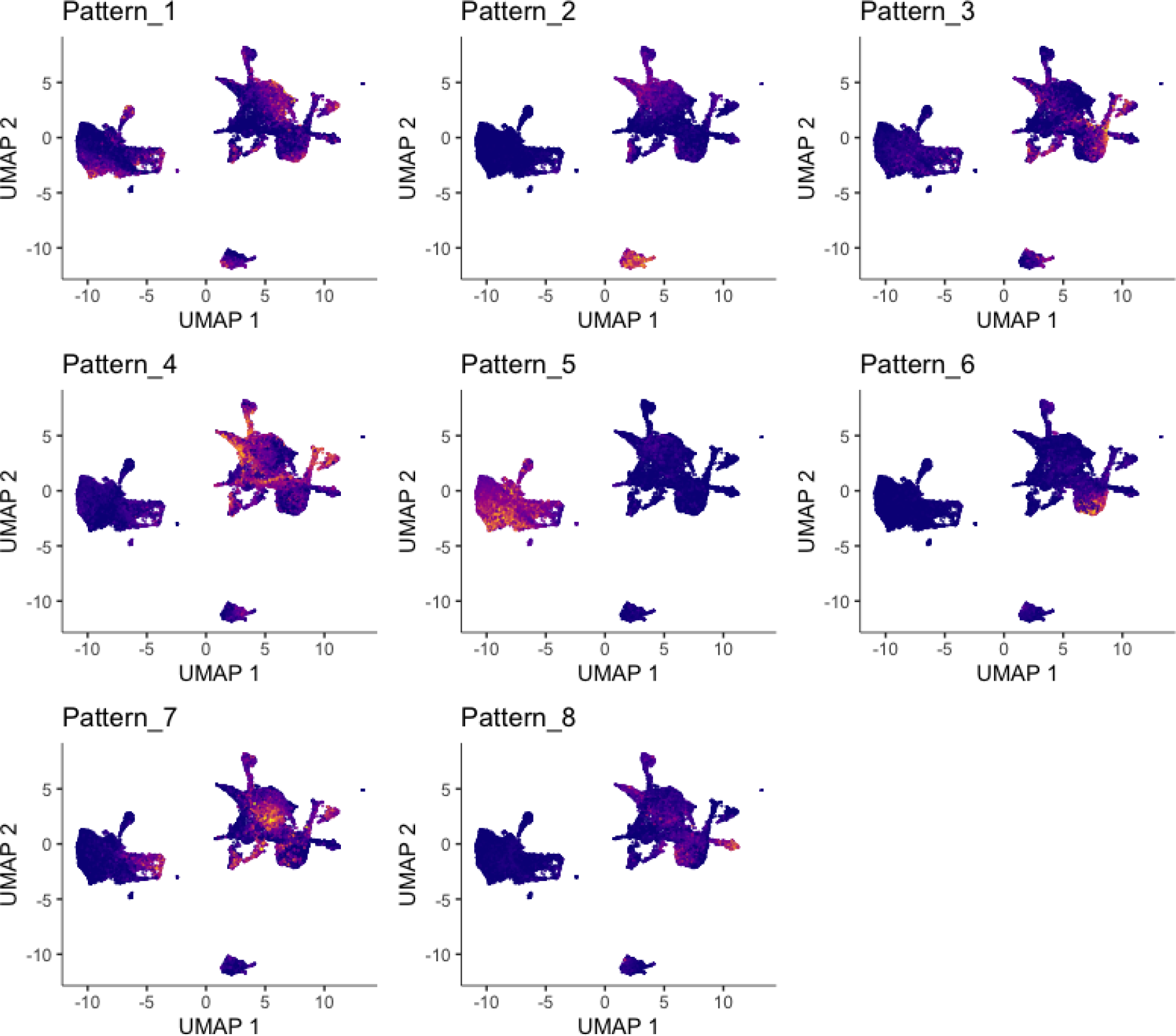

### Assessing Pattern Marker Genes

The patternMarkers() CoGAPS function finds genes associated with each pattern and returns a dictionary of information containing lists of marker genes, their ranking, and their “score” for each pattern. This is vital because genes are often associated with multiple patterns.

patternMarkers can run in two modes, depending on the “threshold” parameter

If **threshold=“all”**, each gene is treated as a marker of one pattern (whichever it is most strongly associated with). The number of marker genes will always equal the number of input genes. If **threshold=“cut”**, a gene is considered a marker of a pattern if and only if it is less significant to at least one other pattern. Counterintuitively, this results in much shorter lists of patternMarkers and is a more convenient statistic to use when functionally annotating patterns.

~~~
pm <- patternMarkers(cogaps, threshold=“cut”)
~~~

### Gene set enrichment analysis

One way to explore and use CoGAPS patterns is by functional annotation of the genes which are significant for each pattern. We have added a method that finds a list of marker genes for each pattern and associates them with msigDB hallmark pathway annotations.

To perform gene set analysis on pattern markers, please run:

~~~
hallmarks <- PatternHallmarks(cogaps)
~~~

hallmarks is a list of data frames, each corresponding to one pattern. To generate a histogram of the most significant hallmarks for any given pattern, please run:

~~~
pl_pattern5 <- plotPatternHallmarks(hallmarks, whichpattern = 5)
pl_pattern5
~~~

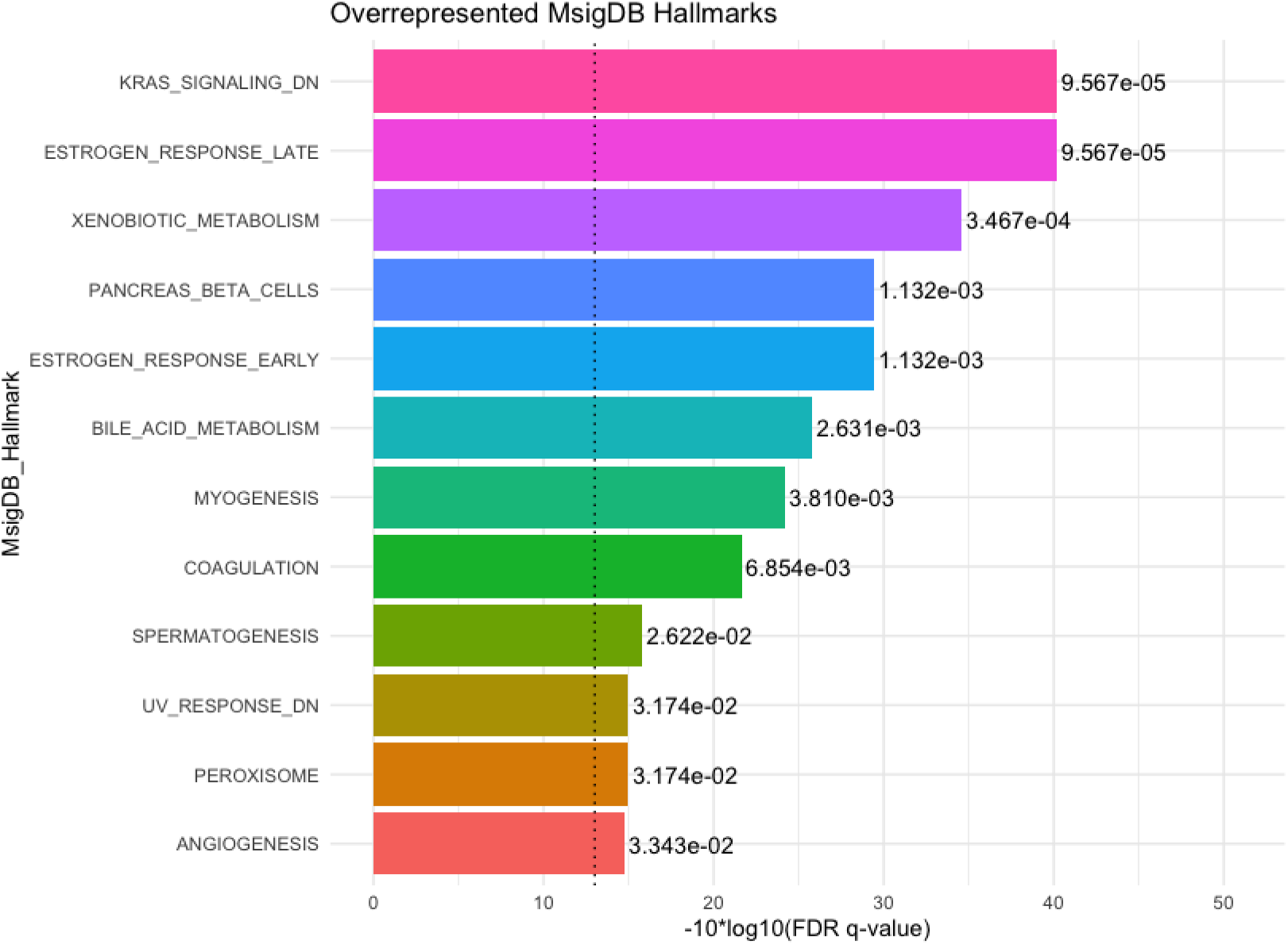

~~~
pl_pattern7 <- plotPatternHallmarks(hallmarks, whichpattern = 7)
pl_pattern7
~~~

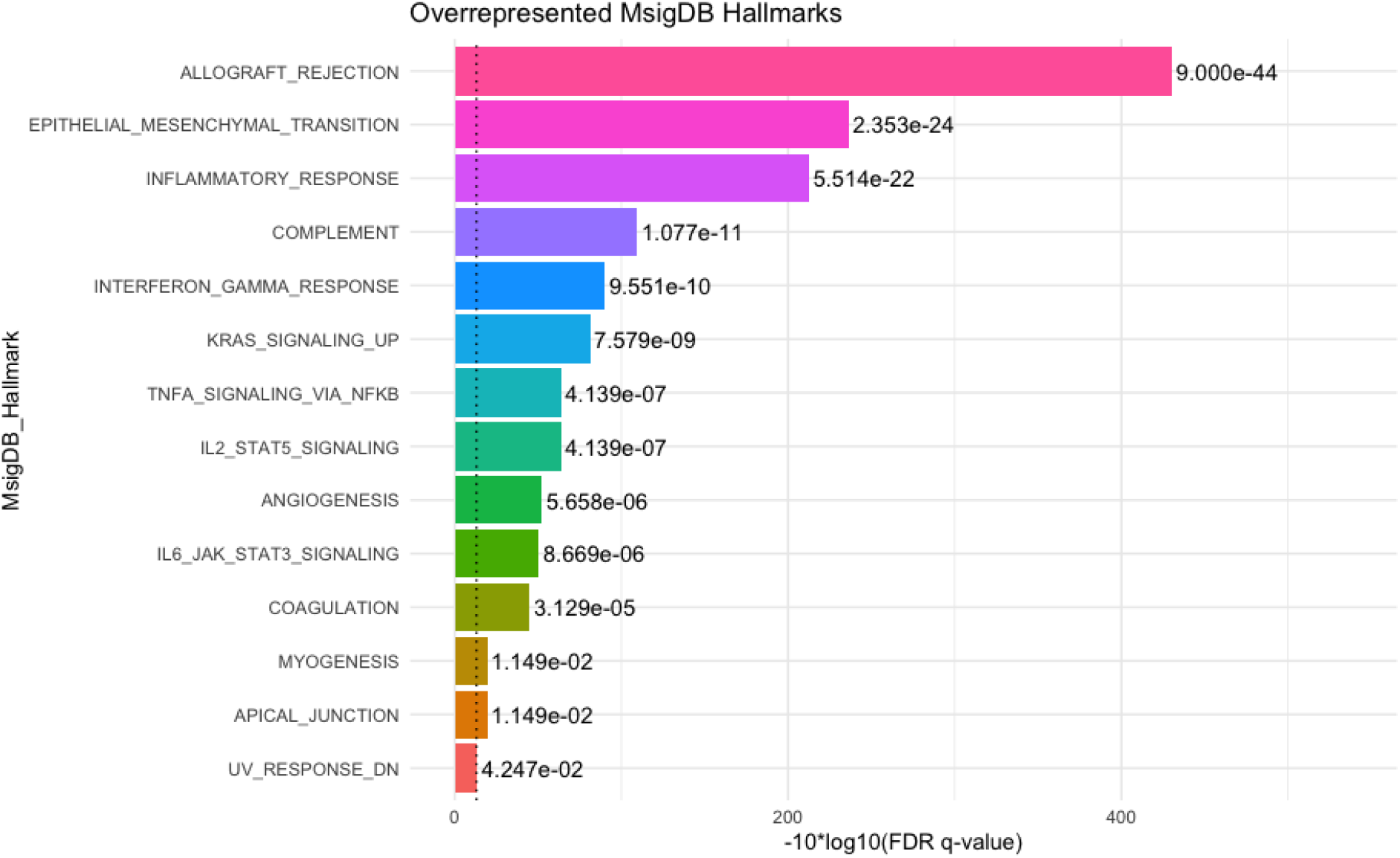

Pattern 5, representing healthy epithelial cell function, has a strikingly different signature of hallmarks than pattern 7.We hypothesize that pattern 7 represents an inflammatory, transitional epithelial cell phenotype as opposed to one centered on tissue maintenance. KRAS signaling is in complete opposition between the two patterns--which makes sense because KRAS is known to be a driving mutation in PDAC.

## Gene Pattern Notebook

The GenePattern Notebook implementation of PyCoGAPS requires no installation of dependencies or local hardware setup. This is highly recommended for users with little to no programming experience, as it provides an interactive, step-by-step walkthrough of running PyCoGAPS. Additionally, GenePattern is a web-based platform that enables users to leverage AWS cloud computing services.

### Steps

1. Log in to the GenePattern Notebook workspace, http://notebook.genepattern.org. If you do not have an account, click the “Register a new GenePattern Account” button, provide the registration information, and log in. Registration for GenePattern Notebook is free.
2. Scroll to “Public Library.” You will see a list of available public notebooks.
3. In the “Search Library” box, search “PyCoGAPS.”
4. Select the “Single-Cell Workflow with PyCoGAPS” notebook by clicking anywhere in its description and selecting “Run Notebook”. A copy of the notebook will be saved in your account.
5. The notebook describes each step in this protocol and contains cells that will allow you to input datasets and set parameters. Follow the instructions in each blue panel, providing information where requested.

### Troubleshooting

*Typical timing is as given in n*log(n). If run times are prohibitive w/in reasonable scales of data may result from algorithm overfitting zeros. Recommend filtering the data only to genes that are reasonably expressed. Filtering to a limited subset of genes (e*.*g*., *high variance) may also help*.

### Timing

As discussed earlier, GIST.csv is representative of a small bulk expression dataset (9 samples x 1363 genes), while GSE98638_HCC.TCell.S5063.count.txt (here abbreviated GSE98638) is a larger matrix, intended as a realistic approximation of data from a single-cell omics experiment or clinical trial of a more average size (5063 cells x 23459 genes). As illustrated below, runtime scales significantly with the input dimension. It is also apparent that Python and R perform similarly on lower-dimension data, while Python has a clear advantage of speed for higher-dimension data.

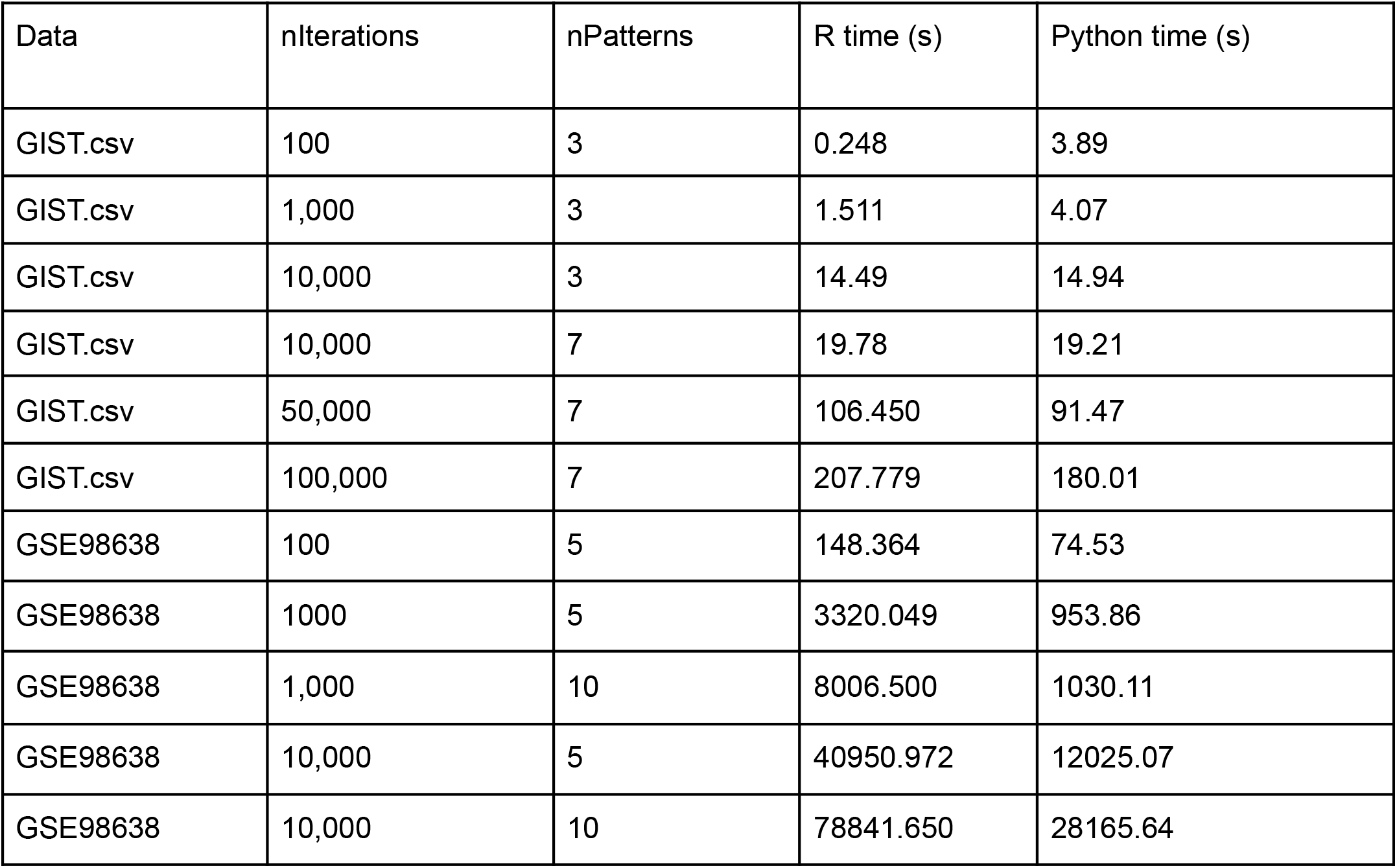

## Data and code availability

The data analyzed in these examples is freely available under accession code GSA: CRA001160, and also from the Genome Sequence Archive, where it has ID: PRJCA001063.

All code is accessible through our lab’s GitHub repositories.

CoGAPS core library and R interface: https://github.com/FertigLab/CoGAPS/

PyCoGAPS (Python interface): https://github.com/FertigLab/pycogaps

## Acknowledgements and Funding

This work was supported by a U24CA248457/U.S. Department of Health & Human Services | National Institutes of Health (J.P.M.), the Chan-Zuckerberg Initiative DAF (2018182718 for Q.H., 2018-183445 to L.A.G., and 2018-183444 to E.J.F.); the Johns Hopkins University Catalyst (E.F. and L.A.G.), an Allegheny Health Network grant (to E.J.F.), U01CA212007 (to E.J.F.), U01CA253403 (to E.J.F.), P01CA247886 (to E.J.F. and E.M.J.), and a Pilot Award from P50CA062924 (to E.J.F.) from the National Cancer Institute, the JHU School of Medicine Synergy Award (to E.J.F. and L.A.G.), 640183 from the Emerson Collective (to E.J.F. and E.M.J), a Kavli Neurodiscovery Institute Distinguished Postdoctoral fellowship (G.S.O.), a Johns Hopkins Provost Award (G.S.O.), K99NS122085 from the BRAIN Initiative in partnership with the National Institute of Neurological Disorders (G.S.O.).

## Contributions

E.J.F., G.S.O., and T.S. originally conceived of the project. E.D.M and M.L. prepared a preliminary draft of the manuscript. A.T. and J.J worked together to write PyCoGAPS, with guidance from G.S.O. A.T. implemented the PyCoGAPS GenePattern Notebook and introduced Docker support. M.R. and J.T.L. provided critical GenePattern Notebook support and collaboration. J.J. and A.T. wrote user guides, and J.T.M. performed the PDAC Atlas single-cell analysis included in them. All authors read, edited, and approved the final manuscript.

