## Supplemental File 1 for "Inferring cellular and molecular processes in single-cell data with non-negative matrix factorization using Python, R, and GenePattern Notebook implementations of CoGAPS"

## This file holds all parameters to be passed into PyCoGAPS.
## To modify default parameters, simply replace parameter values below with user-specified values, and save file. 

# RELATIVE path to data -- make sure to move your data into the created data/ folder
path: data/GIST.csv

# result output file name (output saved as a .pkl file)
result_file: result.pkl

standard_params:
  # number of patterns CoGAPS will learn
  nPatterns: 3
  # number of iterations for each phase of the algorithm
  nIterations: 1000
  # random number generator seed
  seed: 0
  # speeds up performance with sparse data (roughly >80% of data is zero), note this can only be used with the default uncertainty
  useSparseOptimization: False

run_params:
  # maximum number of threads to run on
  nThreads: 1
  # T/F for displaying output
  messages: True
  # number of iterations between each output (set to 0 to disable status updates)
  outputFrequency: 500
  # uncertainty matrix - either a matrix or a supported file type
  uncertainty: null
  # name of the checkpoint file to create
  checkpointOutFile: gaps_checkpoint.out
  # number of iterations between each checkpoint (set to 0 to disable checkpoints)
  checkpointInterval: 250
  # if this is provided, CoGAPS runs from the checkpoint contained in this file
  checkpointInFile: null
  # T/F for transposing data while reading it in - useful for data that is stored as samples x genes since CoGAPS requires data to be genes x samples
  transposeData: False
  # if calling CoGAPS in parallel the worker ID can be specified
  workerID: 1
  # enable asynchronous updating which allows for multi-threaded runs
  asynchronousUpdates: True
  # how many snapshots to take in each phase, setting this to 0 disables snapshots
  nSnapshots: 0
  # which phase to take snapsjots in e.g. "equilibration", "sampling", "all"
  snapshotPhase: sampling

sparsity_params:
  # sparsity parameter for feature matrix
  alphaA: 0.01
  # sparsity parameter for sample matrix
  alphaP: 0.01
  # atomic mass restriction for feature matrix
  maxGibbsMassA: 100
  # atomic mass restriction for sample matrix
  maxGibbsMassP: 100

distributed_params:
  #  either null or genome-wide
  distributed: null
  # number of sets to break data into
  nSets: 4
  # number of branches at which to cut dendrogram used in pattern matching
  # default: nPatterns
  cut: null
  # minimum of individual set contributions a cluster must contain
  # default: math.ceil(cut / 2)
  minNS: null
  # maximum of individual set contributions a cluster can contain
  # default: minNS + nSets
  maxNS: null
  # specify subsets by index or name
  explicitSets: null
  # specify categories along the rows (cols) to use for weighted sampling
  samplingAnnotation: null
  # weights associated with  samplingAnnotation
  samplingWeight: null

additional_params:
  # set of indices to use from the data
  subsetIndices: null
  # which dimension (0=rows, 1=cols) to subset
  subsetDim: 0
  # vector of names of genes in data
  geneNames: null
  # vector of names of samples in data
  sampleNames: null
  # fix either 'A' or 'P' matrix to these values, in the context of distributed CoGAPS, the first phase is skipped and `fixedPatterns: 
  # is used for all sets allowing manual pattern matching, as well as fixed runs of standard CoGAPS
  fixedPatterns: null
  # either 'A' or 'P', indicating which matrix is fixed
  whichMatrixFixed: null
  # whether or not to take PUMP samples
  takePumpSamples: False
  # for reading .h5 files
  hdfKey: null
  # for reading .h5 files
  hdfRowKey: null
  # for reading .h5 files
  hdfColKey: null

aws_params:
  # whether or not to use AWS bucket server
  useAWS: False
  # name of bucket to download from
  downloadBucket: null
  # name of key to download from
  downloadKey: null
  # name of bucket to upload to
  uploadBucket: null
  # name of key to upload to
  uploadKey: null
